# Quantitative imaging of schwannoma captures heterogeneity and accelerates preclinical testing, exposing distinct therapeutic signatures

**DOI:** 10.64898/2026.03.24.714069

**Authors:** Emily A. Wright, Jeremie Vitte, Sara I. Veiga, Sarah E. Bushnell, Catherine E. Movsessian, Youwen Zhang, Jacquelyn Curtis, Ryan B. Corcoran, Shannon L. Stott, Marco Giovannini, Christine Chiasson-MacKenzie, Andrea I. McClatchey

**Affiliations:** Massachusetts General Hospital Krantz Family Center for Cancer Research, Harvard Medical School, 149 13th Street, Charlestown, MA 02129 USA; Department of Pathology, Massachusetts General Hospital, Harvard Medical School, 55 Fruit Street Boston, MA 02114 USA; Department of Head and Neck Surgery, David Geffen School of Medicine at UCLA and Jonsson Comprehensive Cancer Center (JCCC), University of California, Los Angeles, Los Angeles, CA 90095 USA; Department of Medicine, Harvard Medical School, Boston, MA 02115, USA; Center for Engineering in Medicine and Surgery, Massachusetts General Hospital, Boston, MA 02114 USA; The Broad Institute of MIT and Harvard, Cambridge, MA 02142, USA

**Keywords:** schwannoma, *NF2*-schwannomatosis, tumor heterogeneity, preclinical drug testing

## Abstract

Schwannomas are debilitating hallmarks of familial schwannomatoses and common sporadic tumors that form on spinal and cranial nerves. Drug-based therapies for schwannoma are desperately needed but their development has been extremely slow and disappointing, impeded particularly by the poorly understood and surprisingly complex and heterogeneous biology of schwannomas, and by the inefficient use of physiologically relevant *in vivo* preclinical models. We have addressed these gaps by developing a quantitative imaging-centered workflow that allows both a deep analysis of schwannoma development and accelerated preclinical testing in a widely used genetically engineered mouse model of neurofibromatosis type 2-related schwannomatosis (*NF2*-SWN). We deployed our workflow to study schwannoma development and to test three clinically relevant drugs head-to-head. Our results uncovered the very early onset of heterogeneity and macrophage recruitment to initiating schwannomas and unexpectedly revealed that although all three drugs inhibited proliferation within the developing schwannomas, they did so while triggering one of two distinct response signatures, highlighting the value of our pipeline for both rapid drug-testing and translational discovery.

**Significance Statement:** Schwannomas are common sporadic tumors that grow on cranial and spinal nerves, as well as defining features of familial schwannomatosis. Therapeutic advance has been hampered by the striking heterogeneity of schwannomas, Schwann cell plasticity, and the poorly understood biology of interactions between tumor cells and the microenvironment. To address this gap, we exploited a widely used mouse model of schwannomatosis to develop a quantitative imaging platform that captures diverse tumor features, including proliferation, biomarkers of heterogeneity, and macrophage recruitment. We deployed this workflow to holistically compare three clinically active drugs head-to-head and discovered that despite similarly inhibiting proliferation, the drugs induced two distinct response signatures, underscoring the value of this pipeline for accelerated preclinical testing and discovery-based studies.

## Introduction

Familial schwannomatoses are rare tumor predisposition syndromes that feature the development of multiple Schwann cell tumors or schwannomas that predominantly form on and around cranial and spinal nerves, causing loss of hearing, facial paralysis, motor dysfunction and debilitating pain^1,2^. The development of drug-based therapies for schwannoma has been slow and disappointing, yielding largely modest, nondurable responses in a subset of tumors^3–8^. Instead, high-risk surgeries remain the standard of care, after which tumors often recur. As such, new therapies for schwannomatosis patients are urgently needed.

Among many challenges facing therapeutic development for schwannomas is their surprisingly complex and poorly understood biology, which has contributed to slow and inconsistent preclinical pipelines. Schwannomas are genetically non-complex tumors; in humans inherited and/or somatic biallelic inactivation of the *neurofibromatosis type 2* (*NF2)* tumor suppressor gene drives schwannoma development, and mouse models have demonstrated that homozygous *Nf2* inactivation is sufficient for schwannoma formation^9–11^. However, schwannomas are histologically, clinically and therapeutically heterogeneous^12–14^. Recent studies highlight both intrinsic and extrinsic aspects of schwannoma heterogeneity. *In vitro* and in 3D models, *Nf2*^−/−^ Schwann cells are extremely adaptive and able to self-generate spatially patterned heterogeneity that is corroborated in mouse and human schwannoma tissue^15^. Moreover, scRNAseq studies of human schwannoma identify multiple tumor subpopulations, including those expressing signatures of repair or stressed Schwann cells that may reflect an unresolved injury-like state^16–18^. Consistent with this interpretation, macrophages are variably abundant components of human schwannoma and are known to play important roles in the response to peripheral nerve injury and in pathogenic pain, but how or whether they contribute to schwannomagenesis is not known^14,19^.

Few schwannoma cell lines have been established for *in vitro* studies and established lines were derived using different methods and behave dissimilarly *in vitro* and *in vivo*, with some growing aggressively as xenografts, unlike the typically slow, benign growth pattern of schwannomas^15,20–22^. Preclinical studies that capture the complex interactions among schwannoma cell subpopulations, nerves, macrophages and other components of the microenvironment must be carried out *in vivo*. To this end, genetically engineered mouse (GEM) models in which the *Nf2* gene has been conditionally deleted in the Schwann cell lineage have been generated and found to consistently develop multiple spinal schwannomas in the dorsal root ganglia (DRG)^10,23–25^. Preclinical testing of targeted therapies carried out in these mice were validated in human patients^5–8,20,26,27^. However, those studies focused on tumor volume as an endpoint after long-term treatment of mice and the drugs yielded transient, cytostatic responses in both mice and humans. Given our poor understanding of schwannoma biology, an ideal *in vivo* preclinical pipeline would allow the more rapid and efficient evaluation of innovative therapeutic strategies while also amassing insight into why they are or aren’t effective. We have developed a quantitative imaging pipeline to optimize and accelerate the use of a widely utilized GEM model of *NF2*-SWN as a statistically powered tool for both preclinical studies and fundamental discovery. Deployment of this workflow to test three clinically active therapeutic strategies head-to-head uncovered surprisingly distinct impacts that serve as benchmarks for future preclinical studies.

## Results

### Initiation and progression of schwannoma heterogeneity in Postn-Cre;Nf2^flox/flox^ dorsal root ganglia (DRG)

In the widely used *Postn-Cre;Nf2^flox/flox^* GEM model of *NF2*-SWN, lesions develop within all 60 DRG (30 pairs) by 3 months of age^15,25^. Notably, the majority of spinal and vestibular schwannomas in humans also develop within the complex sensory ganglia milieu^28^. Most studies using this and similar GEM schwannoma models have traditionally relied upon caliper-based measurement of DRG volume^20,29,30^ or histological quantification of tumor area in spinal nerve roots^26,31^ as tumor-response endpoints, typically assessed following extended treatment periods. To better understand why drug responses have been modest, and to accelerate the development and testing of new strategies, we set out to holistically capture the biology and heterogeneity of developing and treated lesions at high resolution. Imaging of *Postn-Cre;Nf2^flox/flox^* DRG arrays stained with a general cell membrane marker, as well as markers of satellite glial cells (SGCs) and neuronal soma, reveals that tumor expansion is easily visualized by the progressive separation of neuronal soma, in contrast to the tightly packed soma of control *Nf2^flox/flox^* DRG (Fig. 1A). Using HALO AI software (Indica Labs) to spatially identify the soma, we detect increased intersoma distance beginning as early as 1 month of age along with a corresponding increase in nuclei within the abnormal intersoma tissue (Fig. 1B; Supplemental Fig. 1A-C). Close inspection revealed that at 1 month of age the increased intersoma distance in *Postn-Cre;Nf2^flox/flox^* DRG is driven by the aberrant accumulation of both SGCs that normally cloak individual soma as a monolayer of 2-3 cells, and supernumerary ‘interstitial cells’ between the soma (Fig. 1A). Further accumulation of SGCs and interstitial cells is evident at 3 and 6 months of age and often accompanied by ‘whorl-like’ structures or ‘tumorlets’ that have been described in humans and GEM models and are well-known features of developing and recurring human schwannomas that likely represent multiple independent lesions initiated by *Nf2* deletion^10,32,33^. By 9 and 12 months of age many large whorls and similar structures are present, dramatically separating the soma and distorting the DRG architecture (Fig. 1A,B; Supplemental Fig. 1D).

**Figure 1.**
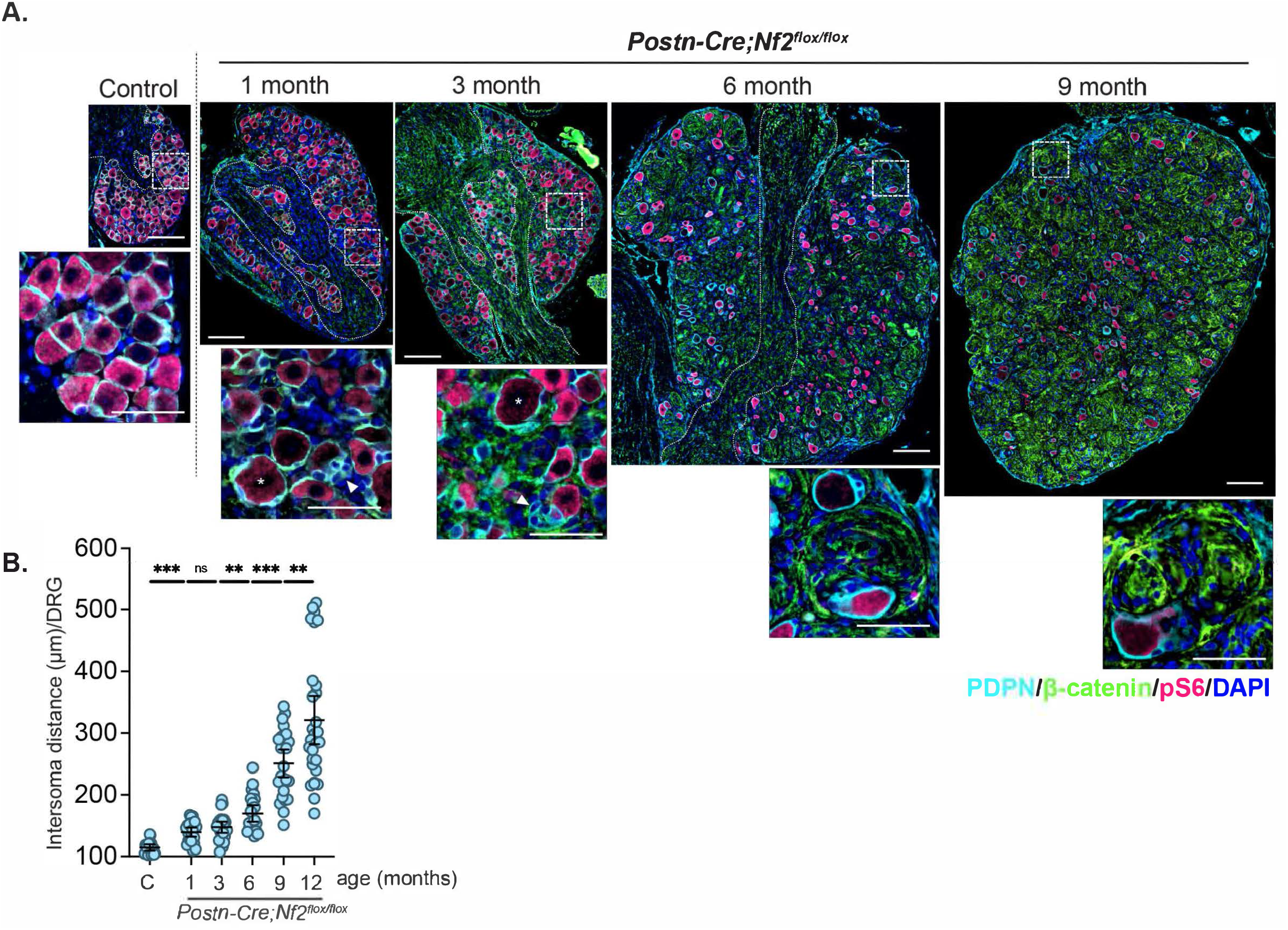
Evolution of schwannomas in *Postn-Cre;Nf2^flox/flox^* DRG. A. Scaled images of representative control (C, 6 month, *Nf2^flox/flox^*^)^ and 1, 3, 6, and 9 month-old mutant (*Postn-Cre;Nf2^flox/flox^*) whole DRG stained with antibodies that detect SGCs (anti-podoplanin (PDPN), cyan) surrounding each neuronal soma, membrane (anti-β-catenin, green), and neuronal soma (anti-pS6; magenta). Nuclei are labeled with DAPI (blue) and the nerve root, where visible in each DRG is demarcated by dotted lines. Insets highlight the cellular organization of aberrant SGCs that form soma-adjacent accumulations (arrowhead) or ‘necklaces’ surrounding individual soma (*), as well as supernumerary interstitial cells that accumulate among the soma and eventually form ‘whorls’ at 6 and 9 months. Note that pS6 also marks some supernumerary interstitial tumor cells. Scale bars: 100 μm main, 50 μm inset. B. Quantitation of distance between soma in control (6 month) and *Postn-Cre;Nf2^flox/flox^* DRG at each timepoint. Each datapoint represents one DRG. Bars represent the mean +/− 95% confidence interval (CI). (**p<0.01, ***p<0.001, ns = not significant).

Macrophages are abundant in human schwannomas and important contributors to the DRG response to nerve injury and other peripheral nerve pathologies^19,34–38^. We found that in contrast to control DRG that have few macrophages at any timepoint, significant numbers of IBA1+ macrophages are present in nascent schwannoma lesions in *Postn-Cre;Nf2^flox/flox^* DRG already at 1 month of age, where many of them are in physical contact with the aberrant SGCs and interstitial cells (Fig. 2A,B; Supplemental Fig. 2A). Macrophages are also present in increased numbers in the nerve root of *Postn-Cre;Nf2^flox/flox^* mice compared to controls at early timepoints (Supplemental Fig. 2B). The number of macrophages then steadily increases and by 6 months of age they often constitute >40% of the cells within each mutant DRG, similar to their abundance in human schwannoma (Fig. 2B; Supplemental Fig. 2C)^38,39^. Notably, throughout the timecourse, similar numbers of macrophages express CD206, a marker of non-inflammatory macrophages, but very few express CD11c, a marker of inflammatory macrophages (Supplemental Fig. 2D-F). Moreover, only a small percentage of cells express the proliferation marker *Top2a*, suggesting that their increase reflects recruitment into the DRG, rather than proliferation of DRG-resident macrophages *in situ* (Supplemental Fig. 2G)^40^. Thus, macrophages may make important and early contributions to schwannoma development.

**Figure 2.**
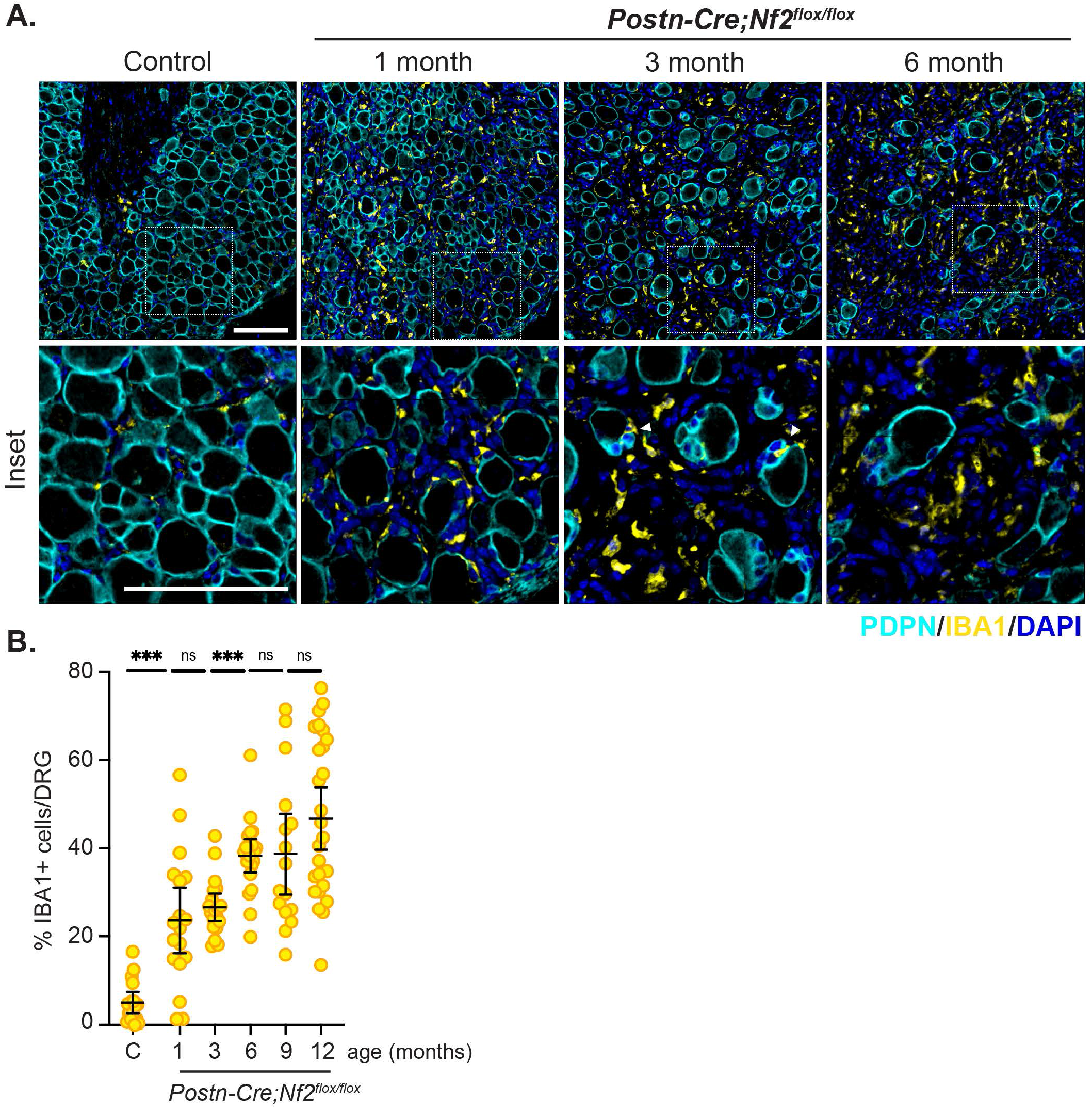
Early and progressive recruitment of macrophages to developing schwannomas in *Postn-Cre;Nf2^flox/flox^* DRG. A. Representative images showing Iba1+ macrophages (yellow) in control (C, 6 month) and *Postn-Cre;Nf2^flox/flox^* DRG at different timepoints. Nuclei are labeled with DAPI (blue). Note that the soma are devoid of staining in these images but are prominently encircled by PDPN+ SGCs. Insets below depict detailed localization of macrophages and their association with both accumulating SGCs (PDPN+; cyan; arrowheads at 3 months) and supernumerary interstitial cells and whorls. Scale bars = 100 μm. B. Quantitation of macrophages in control (6 month) and *Postn-Cre;Nf2^flox/flox^* DRG at several timepoints. Each datapoint represents one DRG. Bars represent the mean +/− 95% CI. (***p<0.001, ns = not significant).

We previously identified phosphorylated S6 ribosomal protein (pS6) and N-myc downstream-regulated gene 1 (pNDRG1) as biomarkers of signaling heterogeneity exhibited by *Nf2^−/−^*Schwann cells *in vitro*, in this GEM model and in human vestibular schwannoma tissue^15^. An examination of how heterogeneity evolves in developing lesions over time revealed quantitatively heterogeneous and dynamic contributions of pS6+ and pNDRG1+ cells (Fig. 3A-C; Supplemental Fig. 3A). Note that pS6 levels are also high in the neuronal soma in both control and mutant DRG, which we excluded from the analysis using a HALO AI tissue classifier (Fig. 1A, 3A). In control DRG at 1 month of age, few cells outside of the soma (masked) are positive for either pS6 or pNDRG1 (Fig. 3A). pNDRG1 levels are normally high in myelinating Schwann cells in the nerve root where it marks the abaxonal, laminin-facing cell compartment (Supplemental Fig. 3A)^41^. However, in mutant DRG many cells in between the soma displayed high levels of pNDRG1 that was notably often localized to the nucleus, which has been linked to a stress-response in other cell types (Fig. 3A,C)^42–45^. Both the distributions and numbers of pNDRG1+ and pS6+ cells in developing lesions were strikingly distinct and heterogeneous, with the total number of pNDRG1+ and pS6+ cells spiking at 1 and 3 months, respectively, and resuming slow, steady increases thereafter, consistent with the small proportion of both populations that express *Top2a* at this timepoint (Fig. 3A,B,D,E; Supplemental Fig. 3A,B). Thus, analysis of each biomarker individually suggests that their distributions are dynamic and spatiotemporally patterned during schwannoma development.

**Figure 3.**
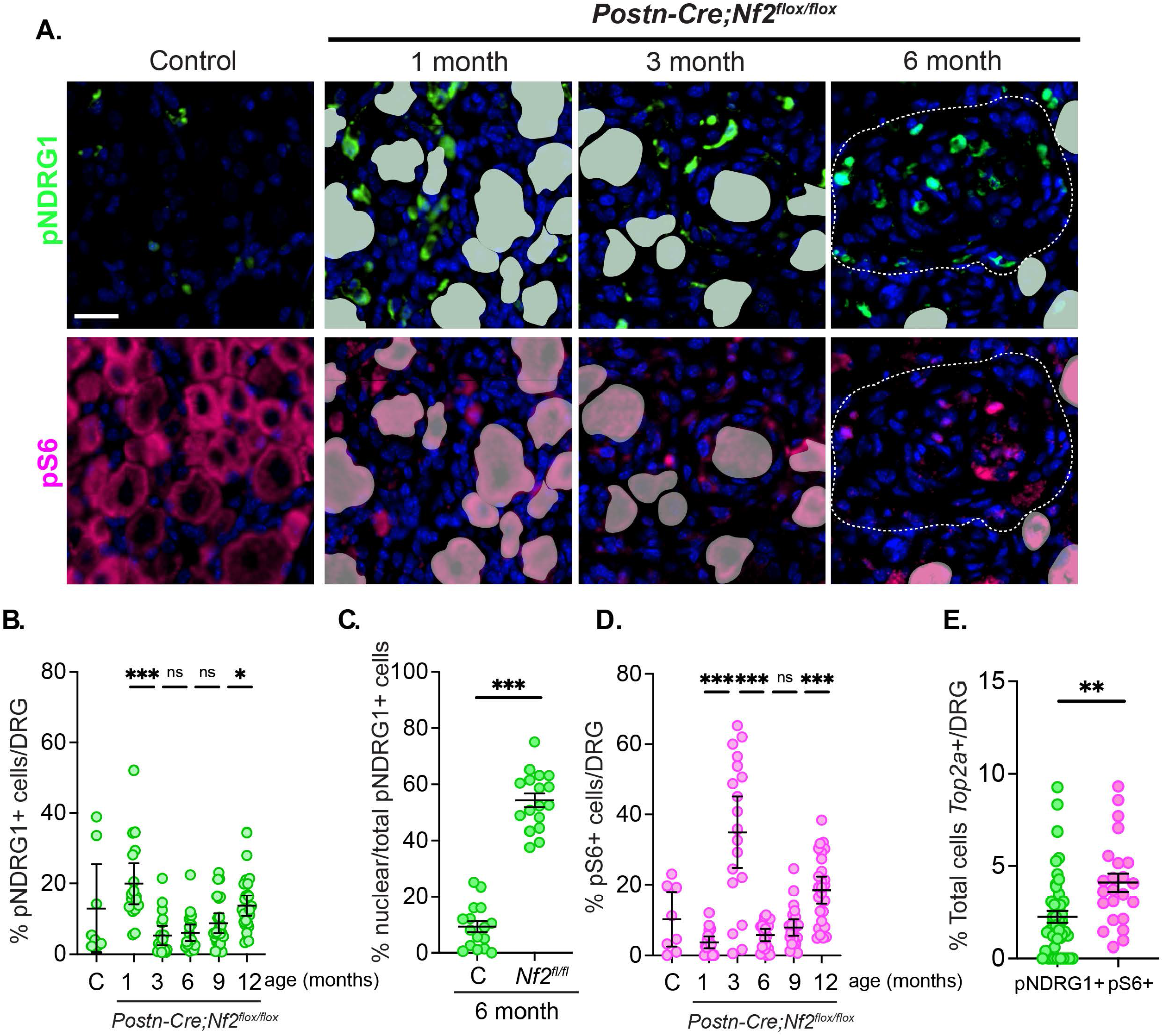
Heterogeneity in developing schwannomas in *Postn-Cre;Nf2^flox/flox^* DRG. A. Representative images of the dynamic and heterogeneous patterns of pNDRG1 (green) and pS6 (magenta) in control (C, 1 month) and *Postn-Cre;Nf2^flox/flox^* DRG at 1, 3, and 6 months of age. Whorl is demarcated by dotted line at 6 months of age. Note that the neuronal soma are pS6+ and masks have been added for visual simplification and orientation; for analyses, tissues were segmented using the Mini Net (HALO AI) classifier to exclude soma from quantitation. Nuclei are labeled with DAPI (blue). Scale bar = 20 μm. B. Quantitation of pNDRG1+ cells in control (6 month) and *Postn-Cre;Nf2^flox/flox^* DRG over time. C. Quantitation of the percentage of pNDRG1+ cells that exhibit nuclear pNDRG1 localization in 6 month-old control and *Postn-Cre;Nf2^flox/flox^* DRG. D. Quantitation of pS6+ cells in control (6 month) and *Postn-Cre;Nf2^flox/flox^* DRG over time. Note that dynamic changes in each biomarker at 1 and 3 months become steady increases by 6 months of age and beyond. E. Quantitation of pNDRG1+ or pS6+ cells that express the proliferation marker *Top2a* as detected by RNAscope in 6 month-old *Postn-Cre;Nf2^flox/flox^*DRG. Each datapoint represents a single DRG. Bars represent the mean +/− 95% CI. (*p<0.05, **p<0.01, ***p<0.001, ns = not significant).

### Quantifying heterogeneity in developing Postn-Cre;Nf2^flox/flox^ schwannomas

To better define the dynamic spatiotemporal patterns of individual biomarkers in developing schwannomas, we used multispectral profiling of individual cells to simultaneously measure SGCs (PDPN+), macrophages (IBA1+), pS6+ and pNDRG1+ cells and identify individual cell populations in tissue arrays of 10-20 DRG (Fig. 4A-C; Supplemental Fig. 4). Multiplex immunofluorescence (mIF) analysis confirmed dynamic changes in pS6 and pNDRG1 in 1 and 3 month-old mutant DRG and also highlighted the appearance of a subpopulation of double positive cells (Fig. 4C). While myelinating pNDRG1+ Schwann cells in the nerve root of control and mutant are rarely also pS6+, there is an emerging subpopulation of double positive interstitial cells in mutant DRG beginning at 3 months of age (Supplemental Fig. 3A; Fig. 4C). Notably, after a period of dynamic onset, the spatiotemporal heterogeneity measured by this analysis stabilized at 6 months of age and was followed by a proportional representation of each biomarker in the markedly expanding tumors (Fig. 2B, 4C). Thus, by 6 months of age schwannomas in *Postn-Cre;Nf2^flox/flox^* DRG exhibit stable intratumoral heterogeneity while steadily increasing in volume.

**Figure 4.**
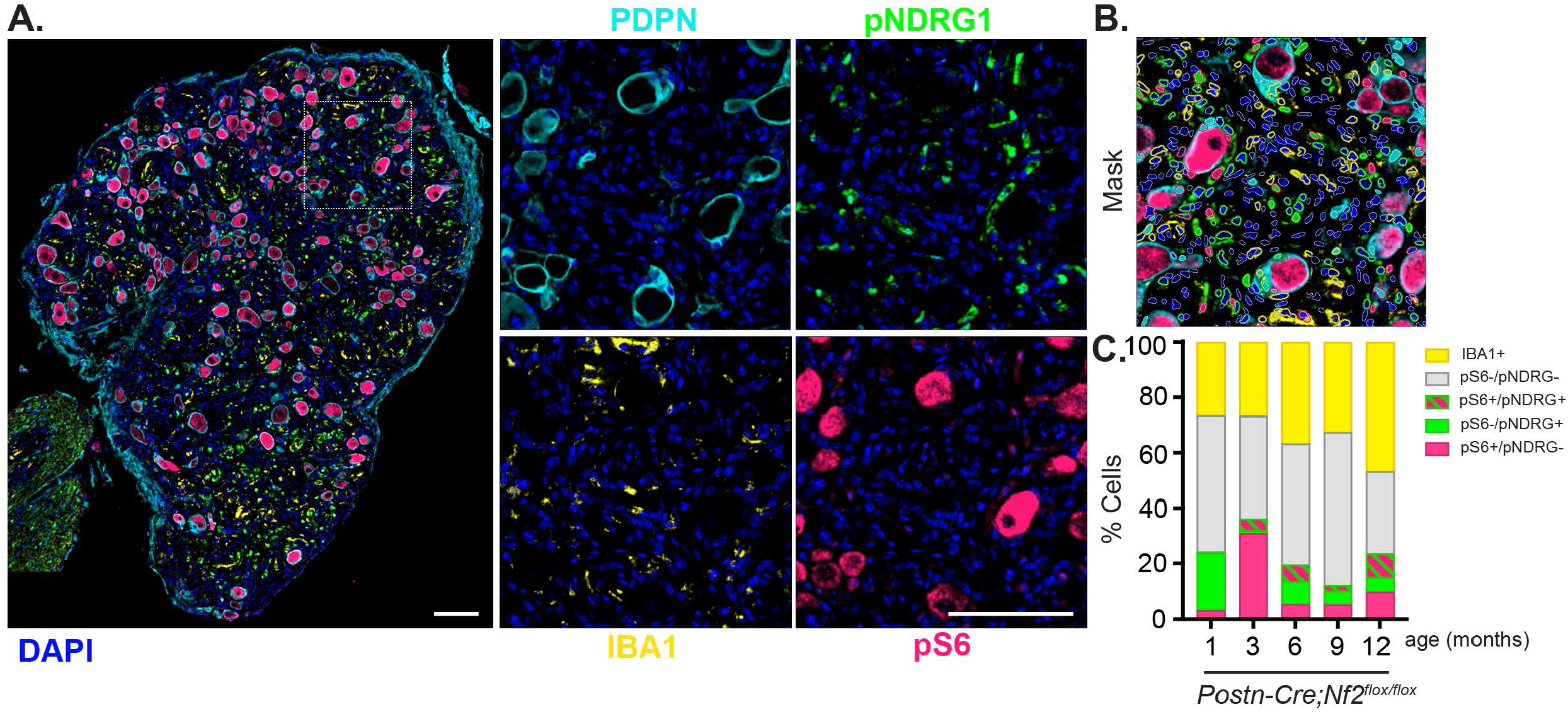
Multiplexed identification of intrinsic and extrinsic markers of heterogeneity in developing schwannomas. A. *Left*, Whole lumbar DRG from a 6 month old *Postn-Cre;Nf2^flox/flox^* mouse stained by mIF for markers of SGCs (PDPN; cyan), macrophages (Iba1; yellow), and schwannoma heterogeneity (pS6; magenta, pNDRG1; green). *Right*, Insets stained for each individual marker. Scale bars = 100 μm. B. Overlay image showing single cell segmentation mask generated using the HighPlex FL module in HALO to identify cells positively stained for each marker. C. Quantitation of individual cell populations over time as percentages of the total number of cells identified. N = 2 mice per timepoint; 6-15 DRG per mouse. Data represent pooled averages of all DRG per timepoint.

### Exploiting multispectral image profiling for quantitative preclinical studies

Optimal preclinical use of the *Postn-Cre;Nf2^flox/flox^*model would exploit the fact that each mouse develops tumors in all 60 DRG^25^. Importantly, in 6 month-old mutant mice measurements of intersoma distance, macrophage numbers, proliferation, and intrinsic tumor biomarker profiles did not uncover differences in lesion formation, expansion or heterogeneity according to either anatomical location (cervical, thoracic or lumbar DRG) or sex (Fig. 5A-B,D-E). Thus, schwannoma formation occurs simultaneously in all 60 DRG of both sexes. Notably, variation in intersoma distance increases at 12 months of age, suggesting the onset of inter-lesion heterogeneity at later timepoints.

**Figure 5.**
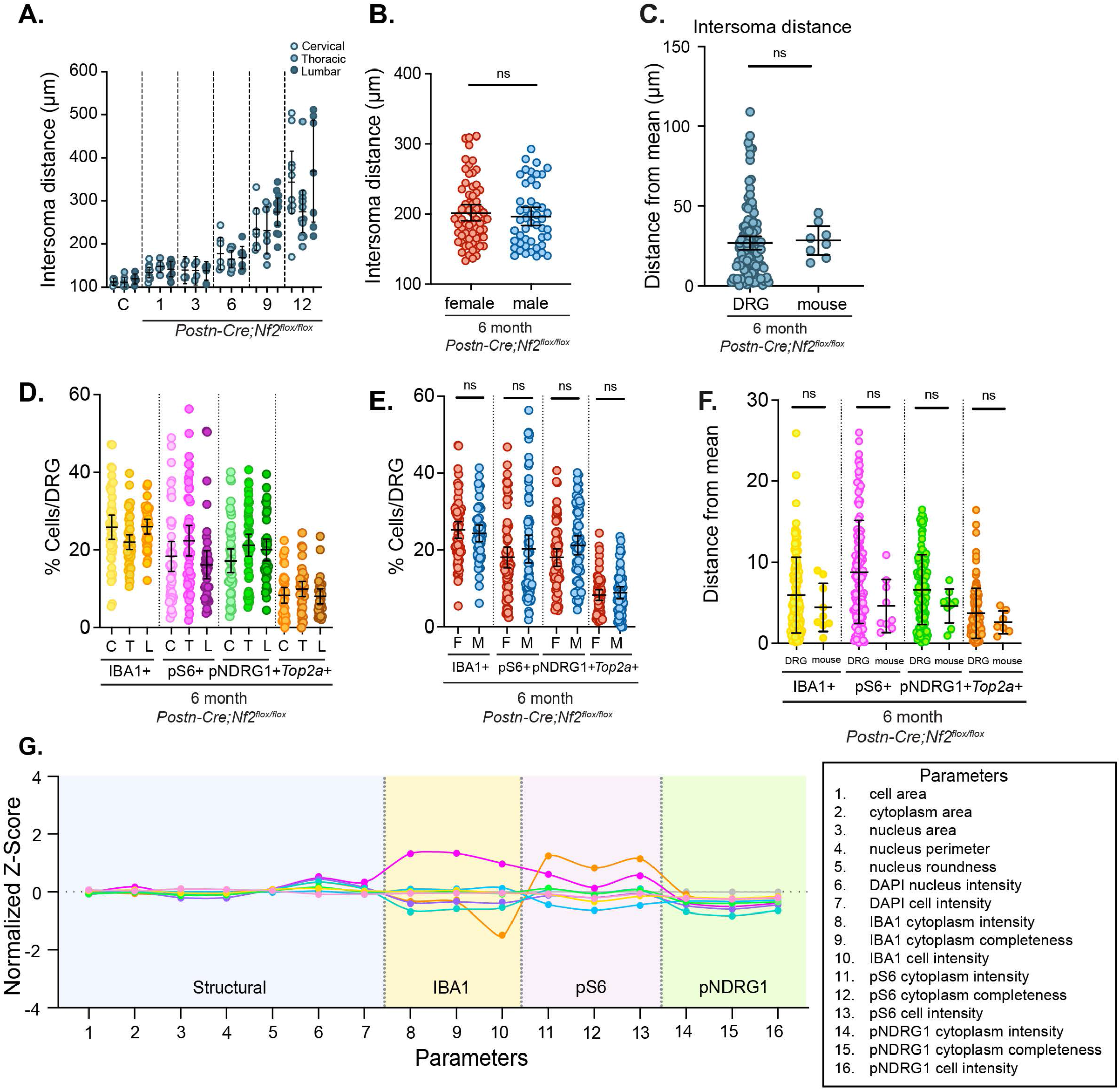
Optimization of *Postn-Cre;Nf2^flox/flox^* DRG analysis for statistically powered preclinical studies. A. Quantitation of intersoma distance according to anatomical location (cervical, thoracic, lumbar) in DRG from 8 *Postn-Cre;Nf2^flox/flox^* mice. Each datapoint represents one DRG. B. Quantitation of intersoma distance according to sex from the same dataset as A. Each datapoint represents one DRG. C. Quantitation of the distance from the mean for DRG-to-DRG and mouse-to-mouse measurements of intersoma distance. For DRG measurements on the left, each data point represents one DRG and for mouse measurements on the right each data point represents the average of 8-23 DRG from each of 8 mice. D. Quantitation of Iba1+ macrophages, pS6+ and pNDRG1+ cells according to anatomical location (C = cervical, T = thoracic, L = lumbar). Each data point represents one DRG. E. Quantitation of Iba1+, pS6+ and pNDRG1+ cells according to sex in DRG from 8 *Postn-Cre;Nf2^flox/flox^* mice. Each data point represents one DRG. F. Quantitation of the distance from the mean for DRG-to-DRG and mouse-to-mouse measurements of Iba1+, pS6+ and pNDRG1+ cells. For DRG measurements on the left, each data point represents one DRG and for mouse measurements on the right each data point represents the average of 8-23 DRG from each of 9 mice. G. Multiparametric profile of multiple DRG from a single 6 month old *Postn-Cre;Nf2^flox/flox^*mouse and parameter legend (right). Parameter numbers are listed on the *x*-axis and normalized Z-scores for each parameter are plotted on the *y*-axis. Parameters are grouped based on similar measurements and are separated by gray dotted lines and shaded accordingly. Each dot represents the normalized Z-score of the mean value of all cells in a DRG for each parameter compared to a reference DRG which was given a Z-score of zero for each parameter (plotted in gray). A continuous line was used to connect adjacent dots.

With this in mind, we compared DRG-to-DRG variation with mouse-to-mouse variation when individual parameters are averaged across DRG (8-23 per mouse) from at least eight 6 month-old *Postn-Cre;Nf2^flox/flox^*mice. We found that DRG-to-DRG and mouse-to-mouse variation were statistically similar for all parameters, justifying the consideration of each DRG as a separate tumor and significantly enhancing the statistical power of our workflow (Fig. 5C,F; Supplemental Fig. 5 A-C). To graphically assemble the results of multiple different tumor features across DRG, we leveraged a multiparametric graphing format to visualize and compare the multiple different measurements for each biomarker reported by HALO AI (Fig. 5G; modified from Collinet et al.)^46^. For example, structural parameters related to cell and nucleus size are measured in multiple ways and are very similar across individual DRG from the same 6 month-old *Postn-Cre;Nf2^flox/flox^* mouse (parameters 1-7), supporting the consistency of our image quantitation. Biological parameters such as macrophages, pS6 and pNDRG1, also measured in multiple ways (parameters 8-10, 11-13, and 14-16, respectively), show more natural variation but remain in good congruence across DRG. Note that the organization of parameter categories in figure 5G was chosen only for visual simplification and can be reordered as more parameters are added. This flexible and scalable multiparametric analysis organizes the immense quantitative content captured by HALO image analysis software, allowing us to profile drug-induced perturbations according to their biological impacts (Fig. 5G).

### Head-to-head multiparametric analysis reveals drug-specific impacts on schwannoma

We deployed our preclinical pipeline to examine two drugs that inhibit clinically active therapeutic targets and have been tested in this or a similar mouse model in long term studies: rapamycin (mTORC1 inhibitor) and brigatinib (FAK/ALK inhibitor)^20,26^. We treated mice for either 7 or 20 days at the same doses used in published studies. For the 20-day treatment, drug was administered on 5 of 7 days per week for a total of 20 days over 4 weeks. Both drugs significantly reduced the number of *Top2a* (83% for rapamycin, 89% for brigatinib) and *Mki67* (97% for rapamycin, 81% for brigatinib) expressing cells after the 20-day treatment, comparable to reductions in tumor volume (75% after 6 weeks of 8 mg/kg rapamycin; 50% after 12 weeks 50 mg/kg brigatinib) or BrdU incorporation (70% after 56 weeks rapamycin treatment) reported after much longer treatments (Fig. 6A; Supplemental Fig. 6A)^20,26^. Importantly, similar reductions occurred after only 7 days of treatment (*Top2a*: 66% rapamycin, 84% brigatinib; *Mki67*: 90% rapamycin, 83% brigatinib). These data suggest that an accelerated 7-day drug treatment may be sufficient to predict long-term drug-induced changes in tumor volume (Fig. 6A; Supplemental Fig. 6A).

**Figure 6.**
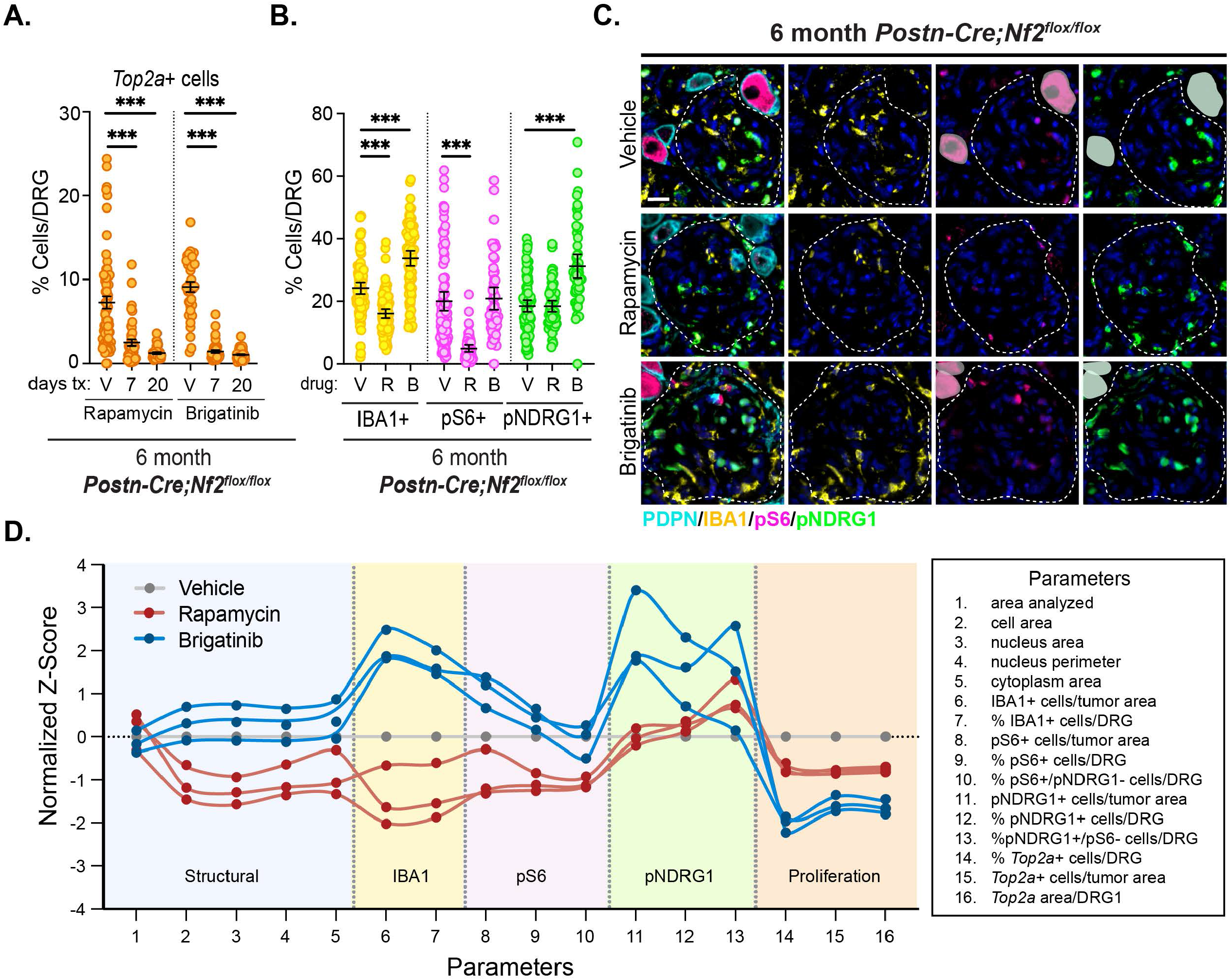
Evaluating two clinically active therapeutic strategies head-to-head. A. Quantitation of *Top2a*+ cells from 6 month-old *Postn-Cre;Nf2^flox/flox^* mice treated for 7 or 20 days (5/7 days per week for 4 weeks) with vehicle (V), rapamycin (R, 8 mg/kg) or brigatinib (B, 50 mg/kg). B. Quantitation of Iba1+, pS6+, and pNDRG1+ cells from 6 month-old *Postn-Cre;Nf2^flox/flox^* mice treated for 7 days with vehicle, rapamycin, or brigatinib. For A and B, each datapoint represents one DRG. Bars represent the mean +/− 95% CI. (***p<0.001). C. Representative images of a *Postn-Cre;Nf2^flox/flox^* DRG showing drug-specific impacts (7-day treatment) on each biomarker in the context of whorls that are demarcated by a dotted line. Soma are masked as in figure 3. Scale bar = 20 μm. D. Multiparametric graph assembling 16 different parameters measured using the HighPlex FL module in HALO across vehicle-(green), rapamycin-(red), or brigatinib-(blue) treated mice (7-day treatment) as described in figure 5. Parameter legend is shown on the right. Each dot represents the normalized Z-score of the mean value of all DRGs from each mouse for each parameter relative to the average values of three vehicle treated mice for each parameter, which was given a Z-score of zero. A continuous line was used to connect adjacent dots for each parameter for each mouse.

Despite the similar impact of each drug on proliferation, their effects on tumor biology were distinct: In addition to eliminating its downstream target pS6 across cell populations without impacting pNDRG1, rapamycin significantly decreased the number of macrophages in each DRG under both drug schedules (Fig. 6B,C; Supplemental Fig. S6B,C). In contrast, brigatinib *increased* both macrophages and pNDRG1+ cells, as well as a further increase in nuclear pNDRG1, while having no impact on pS6 (Fig. 6B,C; Supplemental Fig. S6D). The distinct profiles of change caused by the two drugs are highlighted by multiparametric graphing as well as by principal component analysis (PCA; Fig. 6D; Supplemental Fig. 6E,F,G). Note that rapamycin, but not brigatinib, also led to a significant reduction in cell size, as expected from published studies (Fig. 6D; Supplemental Fig. S6E,F)^26,47^. These data suggest that although rapamycin and brigatinib similarly decrease the growth of schwannomas, they impact schwannoma biology in distinct ways.

Finally, to begin to understand whether therapy-induced phenotypic signatures captured by our holistic pipeline will be unique to each drug tested, we treated a cohort of mice with neratinib, a pan-ErbB inhibitor that is currently being evaluated in NF2 patients (NCT04374305)^7^. Neratinib reduced *Top2a* expression comparably to that of both rapamycin and brigatinib after 7 days of treatment (Fig. 7A; Supplemental Fig. 7). Strikingly however, we found that neratinib provoked a phenotypic signature that was very similar to that of brigatinib and distinct from that of rapamycin (Fig. 7B). Thus, like brigatinib, neratinib treatment increased pNDRG1 levels and macrophage numbers, but did not impact pS6 levels (Fig. 7C). This suggests that therapeutic strategies may elicit discrete classes of biological response in schwannoma and predicts that the clinical response to neratinib in NF2 patients may be similar to that of brigatinib^7^. Altogether, our work establishes an accelerated preclinical pipeline that can holistically capture and classify the differential impacts of clinically relevant drugs on schwannomas in head-to-head evaluation.

**Figure 7.**
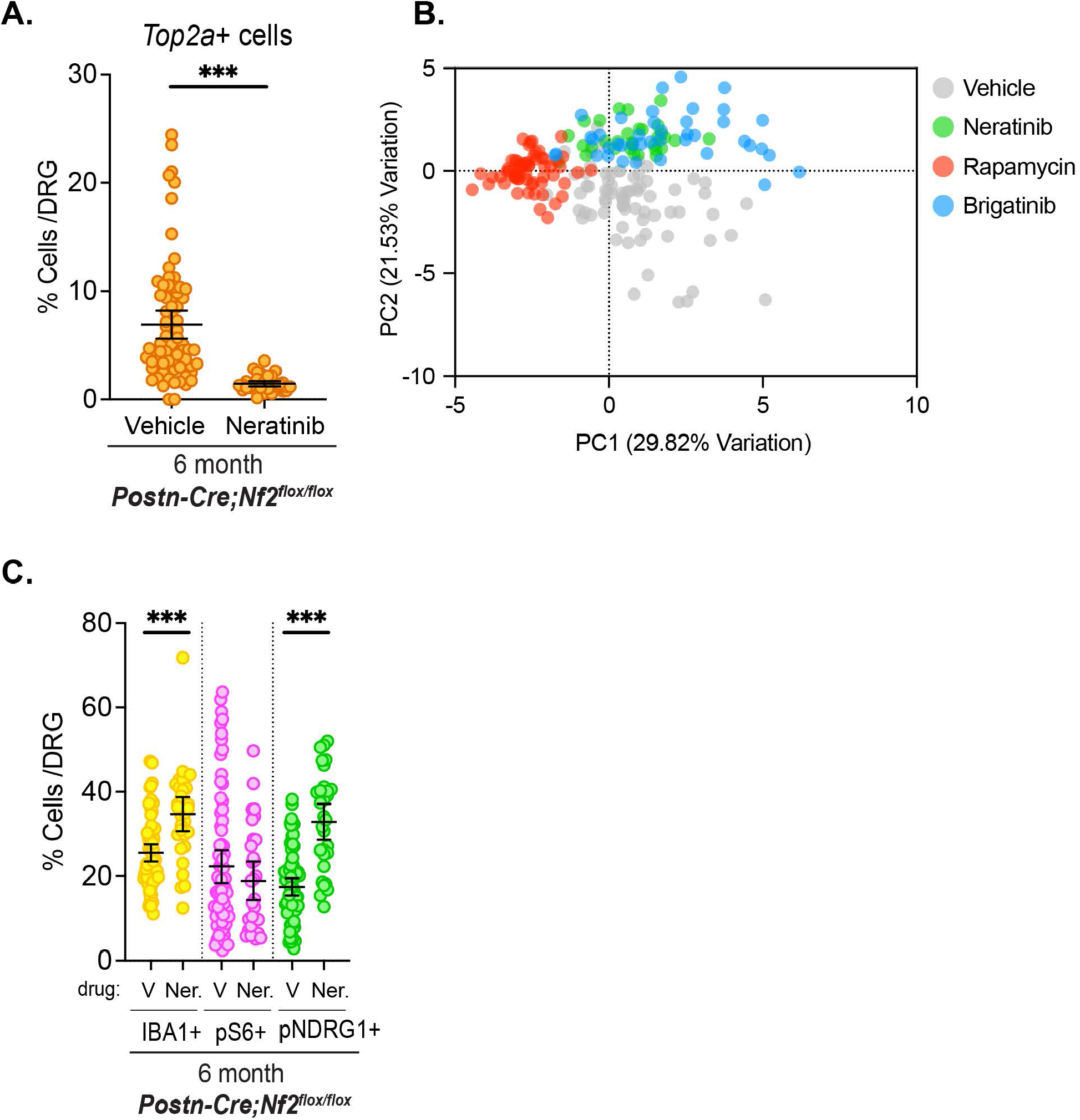
Similarities between brigatinib and neratinib suggest grouped drug responses. A. Quantitation of *Top2a*+ cells from 6 month-old *Postn-Cre;Nf2^flox/flox^* mice treated for 7 days with vehicle (V), or neratinib (N, 40 mg/kg). Each datapoint represents one DRG. Bars represent the mean +/− 95% CI. (***p<0.001). B. Principal component analysis (PCA) plot showing clustering among the parameters measured in figure 6D in brigatinib- and neratinib-treated mice versus those measured in rapamycin- and vehicle-treated mice. Each dot represents one DRG. C. Quantitation of Iba1+, pS6+, and pNDRG1+ cells from 6 month-old *Postn-Cre;Nf2^flox/flox^* mice treated for 7 days with vehicle or neratinib. Each datapoint represents one DRG. Bars represent the mean +/− 95% CI. (***p<0.001).

## Discussion

Familial schwannomatosis patients develop multiple schwannomas of the cranium and spinal cord that are largely treated by surgical removal and/or radiation when they reach debilitating symptomatic thresholds^48^. Few drug-based therapies have been developed and those that have been evaluated provide modest cytostatic relief for growing tumors. Regardless of the treatment, most tumors recur and new ones often form. Thus, a diagnosis of schwannomatosis imposes a lifelong risk of debilitating physical and emotional duress. The pace of therapeutic advance for these syndromes has been excruciatingly slow, due to their rare incidence, the benign nature of the tumors, paucity of human pre- and post-treatment tissue for analysis, preclinical models that require time- and resource-intensive outcome measures, and limited research resources. The innovative and adaptive INTUITT-NF2-designed clinical trial aims to meet some of these challenges by accelerating the testing of new drugs in humans, including brigatinib and neratinib^7^. However, progress also suffers from the surprisingly complex and poorly understood biology of schwannomas themselves which makes it difficult to understand why they do or do not respond to a given therapy. A preclinical pipeline that both accelerates drug testing and advances our understanding of schwannoma biology is desperately needed to meet this gap.

Schwann cells are extremely adaptive and their ability to demyelinate and remyelinate after nerve injury, while also recruiting macrophages to aid in the repair, enables successful peripheral nerve regeneration^49^. Schwannomas have been reported to harbor features of injured nerves and exhibit surprising intrinsic and extrinsic heterogeneity that likely reflects the inherent adaptive capability of Schwann cells^16,18,19,36,38,39^. Notably, while most studies of the role of Schwann cells in nerve injury focus on the distal site of the injury, a less well understood but equally important response occurs in the ganglia, which is where most schwannomas develop^28^. The *Postn-Cre;Nf2^flox/flox^* and related GEM models capture this biology and the slow-growing nature of human schwannomas, while also providing a highly penetrant, early onset, multi-focal model of tumorigenesis that, if used efficiently, enables rapid statistically powered preclinical studies along with the means to mechanistically dissect drug impact now or in the future. In addition to justifying short-term drug treatments, our quantitative establishment that each of 60 DRG in this mouse model can be considered an independent tumor, indicates that many DRG per mouse can be used or banked for complementary analyses such as scRNAseq, proteomics, or metabolomics. Our studies thus establish a framework for the accelerated multi-purpose use of this model for head-to-head testing of new and innovative therapeutic strategies.

Macrophages are abundant components of human schwannomas (up to 53%)^34,38,39,50^ that are also required for the repair of normal nerves^37^, but whether they actually contribute to schwannoma development is not known. Although several studies have reported increased macrophages in rapidly growing human schwannomas^34,51^, our discovery that brigatinib and neratinib both halt proliferation in growing schwannomas but actually increase macrophage numbers (Fig. 6A, 7C), suggests that macrophages may not drive schwannoma growth, at least at relatively early stages. Given that DRG macrophages also play important roles in neuropathic pain, it will be important to determine whether rapamycin, brigatinib and neratinib have differential impacts on pain in this GEM model and in humans. We found that in *Postn-Cre;Nf2^flox/flox^* DRG macrophages are recruited to lesions as soon as they are detectable at one month of age and accumulate rapidly until 6 months of age when they reach about 50%, as in humans, and plateau. At all timepoints up to 12 months of age, the majority of recruited macrophages express CD206, a marker of macrophages that have important pro-regenerative, anti-inflammatory roles in normal nerve repair but are also thought to be pro-tumorigenic in many malignancies and are often dubbed ‘alternatively activated’ or ‘M2-like’ macrophages^52,53^. Recent scRNAseq studies identify significant populations of ‘alternatively activated/M2-like’ macrophages in human schwannoma among more complex macrophage repertoires^16,17,29,38,39,50,51^, justifying the need for a deeper understanding of the complex functional relationships between macrophages and Schwann cell subtypes in both DRG and schwannoma.

Our discovery that clinically active drugs reduce proliferation within expanding schwannomas but have very different impacts on other aspects of schwannoma biology raises key questions for follow-up. It will be important to determine whether the distinct impacts of the drugs on pS6, pNDRG1 and macrophages reflect transient and reversible signaling changes or cell subpopulation state shifts that could create therapeutic vulnerabilities as in other cancers, including glioma^54^. For example, in normal myelinating Schwann cells pS6 and pNDRG1 reflect polarized juxtamembrane signaling that may adaptively maintain membrane homeostasis; on the other hand, in other cell types nuclear pNDRG1, which is evident in many schwannoma cells, is a known stress response^41,55^. In fact, while nuclear pNDRG1+ cells are markedly elevated by 7 days of brigatinib treatment, they return to baseline after 20 days of treatment, and could reflect a transient stress response that could be exploited therapeutically (Fig. 6D, Supplemental Fig. 6D)^42–45^. Our work creates a foundation for holistically understanding both schwannoma development and drug response by layering additional biomarkers, including from scRNAseq datasets, onto this starting atlas. Indeed, our accelerated pipeline and multiparametric analysis can be scaled to add an infinite number of additional parameters. By comparing drugs head-to-head in an efficient way and measuring how they affect multiple aspects of schwannoma biology we can classify therapeutic strategies according to their impacts.

It is important to consider several limitations of our study. First, the genetics of GEM models of schwannoma do not precisely mimic that of human schwannomatosis patients who inherit a heterozygous germline *NF2* mutation that is followed by stochastic somatic inactivation of the remaining copy. Heterozygous *Nf2*-mutant mice that mimic this situation do not spontaneously develop schwannomas, likely because the rate of somatic loss of the wild-type allele is insufficient^56^. In *Postn-Cre;Nf2^flox/flox^* mice homozygous *Nf2* inactivation occurs in all Schwann cells and their precursors in an otherwise wild-type cellular milieu; therefore, each DRG likely harbors many independently initiated ‘lesions’. Indeed, the ‘whorls’ that form in *Postn-Cre;Nf2^flox/flox^*DRG closely resemble ‘tumorlets’ and ‘grape-like’ tumor clusters that are often seen in schwannomatosis patients and thought to reflect independent loss of heterozygosity events^32,33^.

Second, it will be important to determine whether cranial and especially vestibular schwannomas exhibit the same profiles of heterogeneity and drug response that we measured in the DRG models of spinal schwannoma. It has been reported that some *Postn-Cre;Nf2^flox/flox^* mice also develop schwannomas on the 5^th^ and 8^th^ cranial nerves and that some suffer loss of hearing^25^. Although brigatinib more significantly impacted non-vestibular schwannomas in humans^7^, it has recently been reported that knockout or pharmacologic inhibition of one of the key brigatinib targets, FAK, both reduces DRG volume and improves hearing in *Postn-Cre;Nf2^flox/flox^* mice^29^. However, cranial nerve tumors in these mice are less tractable, developing with less predictable penetrance and onset, and much lower multiplicity^25^. The quantitative and biological analysis of spinal tumors in these mice will provide a crucial framework to guide the future study of those tumors.

Finally, we focused on reduced proliferation as a primary endpoint, intentionally mirroring previous preclinical studies that measured volumetric changes in tumor growth, which will not capture other functional endpoints such as improved pain, motor function, or hearing^20,26,29^. However, follow-up studies that link drug-response profiles identified by our platform to other endpoints could yield valuable predictive and mechanistic information. Finally, although our quantitative imaging analyses together with analyses of banked DRG from treated mice will capture troves of additional biological information that can be deeply analyzed by comparison to other drugs as they are tested, short-term drug treatments may not capture longer-term impacts such as cell state changes that may also impact tumor biology and the emergence of drug resistance. The comparative profiles of short-term responses to multiple drugs captured by our pipeline can serve as a basis for selecting certain drugs for follow up studies using other endpoints, longer term treatments and drug combinations.

## Materials and Methods

### Mice

The *Postn-Cre;Nf2^flox/flox^*genetically engineered mouse model of *NF2*-SWN was generated by breeding *Postn-Cre* and *Nf2^flox^*mice as previously described^15,23,25^. All animal care and experimentation were performed with the approval of the Mass General Brigham and UCLA Institutional Animal Care and Use Committees. Genotypes of all offspring were confirmed by PCR analysis using genomic DNA obtained from tail biopsies using the following primers: The *Nf2^flox^* allele was detected using primers F (5′-CTTCCCAGACAAGCAGGGTTC-3′) and R (5′-GAAGGCAGCTTCCTTAAGTC-3′), yielding a 442-bp (*Nf2^flox^* allele) and 305-bp (*Nf2^WT^* allele) product. The *Postn-Cre* transgene was detected using primers cre-s1 (5′ACATGTTCAGGGATCGCCAG-3′) and cre-a1 (5′-TAACCAGTGAAACAGCATTGC-3′), yielding a 230-bp product. Mice were housed under standard conditions, monitored bi-weekly and euthanized by CO_2_ inhalation prior to tissue harvest.

### *In vivo* studies

Beginning at 6 months of age, *Postn-Cre;Nf2^flox/flox^*mice were treated with rapamycin (MedChemExpress, Cat# HY-10219), brigatinib (MedChemExpress, Cat# HY-12857) or neratinib (MedChemExpress Cat # HY-32721). For short-term treatments, drugs were administered once daily for 7 consecutive days, and for long-term treatments, drugs were administered once daily on 5 of 7 days per week for a total of 20 days over 4 weeks. Rapamycin was administered by intraperitoneal (IP) injection at 8 mg/kg in vehicle (95.5% H₂O, 4% DMSO, 0.25% polyethylene glycol 400, and 0.25% Tween-80); brigatinib was administered by oral gavage at 50 mg/kg in vehicle (25 mM sodium citrate buffer pH 4.5); and neratinib was administered by oral gavage at 40 mg/kg in vehicle (50% saline, 44.4% polyethylene glycol 400, and 5.6% tween 80). Animals were monitored daily for signs of distress, including lethargy, decreased mobility or hunched posture. Mice were euthanized and DRG were harvested 1 hour after the final administration of drug.

### Tissue preparation

Immediately following humane euthanasia, DRG were microscopically dissected following standard protocols and categorized according to anatomical location (cervical, thoracic, lumbar). Tissues were fixed in 10% neutral buffered formalin for 24 hours before dehydrating in graded ethanol solutions, clearing with xylene, infiltrating with molten paraffin, and embedding in paraffin blocks as tissue arrays of 10-20 DRG organized by anatomical location. Tissue arrays were constructed based on anatomical location. Formalin-fixed paraffin-embedded (FFPE) blocks were cut into 5 μm sections and deparaffinized in xylene and rehydrated through graded ethanol solutions prior to staining.

### Multiplex Immunofluorescence (mIF)

Multiplex immunofluorescence (mIF) staining was performed on FFPE DRG sections using the Opal-based tyramide signal amplification system (Quanterix). Briefly, DRG sections were subject to heat-induced antigen retrieval in sodium citrate buffer pH 6.0 (AR600250ML, Quanterix), incubated in blocking solution (ARD1001EA, Quanterix) for 10 minutes, and stained with primary antibodies diluted in antibody diluent (S3022, DAKO) for 30 minutes at room temperature. Sections were then incubated with species specific HRP-conjugated secondary antibodies followed by signal amplification with Opal fluorophores (see Antibodies section below, Quanterix) diluted in 1X Plus Amplification Diluent (FP1609, Quanterix) for 10 minutes. Antibody complexes were removed between staining cycles using heat-mediated stripping in sodium citrate buffer pH 6.0, allowing sequential detection of multiple targets. Nuclei were counterstained with 4’,6-diamidino-2-phenylindole (DAPI), and coverslips were mounted with ProLong gold (Invitrogen) mounting medium and allowed to cure overnight.

### RNA-scope/in situ hybridization (ISH)

FFPE tissue sections were stained following the Integrated Co-detection Workflow combining ISH using the RNAscope Multiplex Fluorescent v2 assay (ACD, Advanced Cell Diagnostics) and immunofluorescence as described^14^. After tissue preparation according to ACD recommendations (Protocols 323100-USM and MK 51-150/Rev B), sections were steamed for target retrieval in sodium citrate pH 6.0 (15 minutes). Tissues were incubated with primary antibodies overnight at +4°C. Hybridization was performed using probes targeting mouse *Top2a* (Cat#491221, ACD) or *Mki67* (Cat#416771 or Cat#416771-C2, ACD). Opal fluorophores were used for signal development. Primary antibodies were detected with rabbit HRP-conjugated secondary antibodies followed by amplification with opal fluorophores. Nuclei were counterstained with DAPI, and slides were mounted with ProLong gold.

### Multispectral Imaging and Analysis

All slides were imaged on the Vectra3 quantitative pathology imaging system, using the 40x objective, NA 0.75 (Quanterix). A custom spectral library was generated in InForm 2.5.1 as previously described using the Schwann cell marker SCN7A to generate library slides stained with each fluorophore^57^. For each DRG section, the entire tissue area was scanned on the Vectra3, annotated in Phenochart, and raw images spectrally unmixed in InForm for import into our data analysis platform (HALO AI v4.0.5107.488 Indica Labs). Unmixed images were fused using the HALO software. For each slide, annotation layers were manually traced to select DRG tissue and exclude nerve root and a classifier was created using the MiniNet (HALO AI) module to exclude neuronal cell bodies from antibody and mRNA analysis. Single-cell segmentation and quantitation was performed using the HighPlex FL v4.3.2. and FISH-IF v2.3.1 HALO AI modules respectively. For intersoma distance calculations, neuronal cell bodies were identified and centroid coordinate data was obtained using the Nuclei Seg (HALO AI) classifier.

### Antibodies

The following primary antibodies were used: Anti-Podoplanin hamster monoclonal antibody (mAb) (1:5000; clone eBio8.1.1, 14-5381-85, Invitrogen), anti-pS6 (S235/236) rabbit polyclonal antibody (pAb) (1:5000; 2211, Cell Signaling Technology), anti-pNDRG1 (T346) rabbit mAb (1:5000; clone D98G11, 5482, Cell Signaling Technology), anti-IBA1 rabbit pAb (1:2000; GTX100042, GeneTex), anti-β-catenin rabbit mAb (1:5000; clone E247, AB32572, Abcam), anti-CD206/MRC1 rabbit mAb (1:2000; clone E6T5J, 24595 Cell Signaling Technology), anti-CD11c rabbit mAb (1:2000; clone D1V9Y, 97585, Cell Signaling Technology), antiSCN7A rabbit pAb (1:5000; NB100-81029, Novus Biologicals), anti-pEGFR (Y1068) rabbit mAb (1:1000; clone D7A5, 3777, Cell Signaling Technology), and anti-EGFR rabbit pAb (1:1000, 06-847, Millipore Sigma. The following secondary antibodies were used for mIF: Polink-2 plus HRP Syrian Hamster DAB (3,3’-Diaminobenzidine) Detection Kit (D86-18, OriGene) and Polink-2 plus HRP Rabbit DAB Detection Kit (D39-18, OriGene). The following fluorophores were used (all from Quanterix): Opal 520 Reagent Pack (1:300; FP1487001KT), Opal 570 Reagent Pack (1:300; FP1488001KT), Opal 620 Reagent Pack (1:300; FP1495001KT), and Opal 650 Reagent Pack (1:300; FP1496001KT). HRP-conjugated secondary anti-mouse and anti-rabbit antibodies for western blotting were from Amersham.

### Western blotting

Cell pellets were lysed in mRIPA buffer with protease inhibitors (1% Triton X-100, 0.5% sodium deoxycholate, 0.1% SDS, 50 mM Tris-HCl at pH 7.4, 140 mM NaCl, 1 mM EDTA, 1 mM EGTA, 1 mM PMSF, 1 mM Na_3_VO_4_, 10 µg/mL leupeptin, 10 µg/mL pepstatin, 10 µg/mL aprotinin), for 1 h on ice, then lysates cleared by centrifugation. 40 µg of protein were loaded on SDS-PAGE gels, transferred to PVDF membranes, and incubated at 4° C overnight with antibodies diluted in 5% milk or 5% BSA (for phospho-specific antibodies). The membranes were then incubated with secondary antibodies, exposed to ECL substrate and film-developed.

### Multiparametric Analysis

We adapted our multiparametric analysis pipeline from Collinet et al^46^. We identified a set of values measured by the HighPlex FL v4.3.2. HALO AI module that captured features of cellular structures and individual biomarkers and together define the phenotypic profile of each DRG. These parameters were categorized into structural features, IBA1+ cell features, pS6+ cell features, pNDRG1+ cell features, and proliferating cell features. For DRG-to-DRG comparisons, object data from individual cells for each parameter was collected, a random DRG was chosen as a reference profile, and data for each parameter for each DRG was normalized according to the mean and standard deviation of the reference data set to obtain a normalized Z-score using the formula:

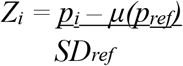

where *p_i_* equals the object data of a parameter, *μ(p_ref_)* equals the mean value of each parameter in the reference DRG and *SD_ref_* equals the standard deviation of each parameter in the reference DRG. For mouse-to-mouse comparisons in the drug treatment experiments, summary data for individual parameters for each DRG were extracted from HALO and normalized according to the mean and standard deviation of a reference dataset composed of the average of vehicle-treated mice for each cohort (n=3). Z-values were calculated using the above formula. PCA was performed in Prism11 (Graphpad Software) using standardized individual DRG values of the same parameters used for multiparametric analysis, with the number of components retained determined by parallel analysis (1000 simulations, 95% confidence interval).

### Statistics and Reproducibility

Data from all analyses were imported into Prism10 for plotting graphs and statistical analysis. The unpaired two tailed Welch’s t test was used to compare differences between two groups. One way ANOVA with Tukey’s multiple comparisons test was used to compare data across 3 or more groups. All data are representative of at least three independent experiments unless otherwise specified.

## Data Availability

The data that support the findings of this study are available in the article or the supplemental information files.

## Acknowledgments.

We would like to thank present and past members of the McClatchey laboratory and Krantz Family Center for Cancer Research for valuable discussions. Harvard Cancer Consortium in Boston, MA, for the use of the MGH Specialized Histopathology Core, which provided tissue processing, embedding, and cutting services. Harvard Cancer Consortium is supported in part by an NCI Cancer Center Support Grant # NIH 5 P30 CA06516. This work was supported by the U.S. Army Medical Research and Development Command, through the Neurofibromatosis Research Program under Award Nos. W81XWH-19-0156 (A.I.M.), W81XWH-21-1-0446 (A.I.M.), W81XWH-16-10086 (M.G.), and W81XWH-21-1-0448 (M.G.), and by Children’s Tumor Foundation Young Investigator Award 2023-01-007 (S.I.V.)

## Author Contributions

The study was conceived and designed by C.C.M., J.V., M.G., and A.I.M. In vivo experiments were carried out by E.A.W., C.C.M., J.V., S.I.V, S.E.B., C.E.M., Y.Z., and J.C. Whole-slide scanning and HALO image analysis were carried out by E.A.W., C.C.M., S.I.V., S.E.B., and C.E.M. Data were analyzed and interpreted by E.A.W., C.C.M., S.E.B., C.E.M, J.V., M.G., and A.I.M. The project was supervised by C.C.M., A.I.M., M.G., S.L.S, and R.B.C. Drafting of the manuscript and preparation of figures was completed by E.A.W., C.C.M., J.V., and A.I.M. All authors read and commented on the manuscript.

## References

1. Plotkin, S.R., et al. Updated diagnostic criteria and nomenclature for neurofibromatosis type 2 and schwannomatosis: An international consensus recommendation. Genet Med 24, 1967–1977 (2022).

2. Evans, D.G. Neurofibromatosis type 2. Handb Clin Neurol 132, 87–96 (2015).

3. Karajannis, M.A., et al. Phase II trial of lapatinib in adult and pediatric patients with neurofibromatosis type 2 and progressive vestibular schwannomas. Neuro Oncol 14, 1163–1170 (2012).

4. Blakeley, J.O., et al. Efficacy and Biomarker Study of Bevacizumab for Hearing Loss Resulting From Neurofibromatosis Type 2-Associated Vestibular Schwannomas. J Clin Oncol 34, 1669–1675 (2016).

5. Goutagny, S., et al. Phase II study of mTORC1 inhibition by everolimus in neurofibromatosis type 2 patients with growing vestibular schwannomas. J Neurooncol 122, 313–320 (2015).

6. Goutagny, S., Giovannini, M. & Kalamarides, M. A 4-year phase II study of everolimus in NF2 patients with growing vestibular schwannomas. J Neurooncol 133, 443–445 (2017).

7. Plotkin, S.R., et al. Brigatinib in NF2-Related Schwannomatosis with Progressive Tumors. N Engl J Med 390, 2284–2294 (2024).

8. Nghiemphu, P.L., et al. Imaging as an early biomarker to predict sensitivity to everolimus for progressive NF2-related vestibular schwannoma. J Neurooncol 167, 339–348 (2024).

9. Agnihotri, S., et al. The genomic landscape of schwannoma. Nat Genet 48, 1339–1348 (2016).

10. Vitte, J., Gao, F., Coppola, G., Judkins, A.R. & Giovannini, M. Timing of Smarcb1 and Nf2 inactivation determines schwannoma versus rhabdoid tumor development. Nat Commun 8, 300 (2017).

11. Stemmer-Rachamimov, A.O., et al. Universal absence of merlin, but not other ERM family members, in schwannomas. Am J Pathol 151, 1649–1654 (1997).

12. Erlandson, R.A. & Woodruff, J.M. Peripheral nerve sheath tumors: an electron microscopic study of 43 cases. Cancer 49, 273–287 (1982).

13. Wippold, F.J., 2nd, Lubner, M., Perrin, R.J., Lammle, M. & Perry, A. Neuropathology for the neuroradiologist: Antoni A and Antoni B tissue patterns. AJNR Am J Neuroradiol 28, 1633–1638 (2007).

14. Lewis, D., et al. The microenvironment in sporadic and neurofibromatosis type II-related vestibular schwannoma: the same tumor or different? A comparative imaging and neuropathology study. J Neurosurg 134, 1419–1429 (2020).

15. Chiasson-MacKenzie, C., et al. Cellular mechanisms of heterogeneity in NF2-mutant schwannoma. Nat Commun 14, 1559 (2023).

16. Barrett, T.F., et al. Single-cell multi-omic analysis of the vestibular schwannoma ecosystem uncovers a nerve injury-like state. Nat Commun 15, 478 (2024).

17. Gonzalez Castro, L.N., et al. A single-cell atlas of Schwannoma across genetic backgrounds and anatomic locations. Genome Med 17, 37 (2025).

18. Liu, S.J., et al. Epigenetic reprogramming shapes the cellular landscape of schwannoma. Nat Commun 15, 476 (2024).

19. Hannan, C.J., et al. The inflammatory microenvironment in vestibular schwannoma. Neurooncol Adv 2, vdaa023 (2020).

20. Chang, L.S., et al. Brigatinib causes tumor shrinkage in both NF2-deficient meningioma and schwannoma through inhibition of multiple tyrosine kinases but not ALK. PLoS One 16, e0252048 (2021).

21. Synodos for, N.F.C., et al. Traditional and systems biology based drug discovery for the rare tumor syndrome neurofibromatosis type 2. PLoS One 13, e0197350 (2018).

22. Troutman, S., et al. Crizotinib inhibits NF2-associated schwannoma through inhibition of focal adhesion kinase 1. Oncotarget 7, 54515–54525 (2016).

23. Giovannini, M., et al. Conditional biallelic Nf2 mutation in the mouse promotes manifestations of human neurofibromatosis type 2. Genes Dev 14, 1617–1630 (2000).

24. Giovannini, M., et al. Schwann cell hyperplasia and tumors in transgenic mice expressing a naturally occurring mutant NF2 protein. Genes Dev 13, 978–986 (1999).

25. Gehlhausen, J.R., et al. A murine model of neurofibromatosis type 2 that accurately phenocopies human schwannoma formation. Hum Mol Genet 24, 1–8 (2015).

26. Giovannini, M., et al. mTORC1 inhibition delays growth of neurofibromatosis type 2 schwannoma. Neuro Oncol 16, 493–504 (2014).

27. Karajannis, M.A., et al. Phase II study of everolimus in children and adults with neurofibromatosis type 2 and progressive vestibular schwannomas. Neuro Oncol 16, 292–297 (2014).

28. Tryggvason, G., et al. Radiographic association of schwannomas with sensory ganglia. Otol Neurotol 33, 1276–1282 (2012).

29. Mitchell, D.K., et al. Inhibition of focal adhesion kinase impairs tumor formation and preserves hearing in a murine model of NF2-related schwannomatosis. Sci Adv 12, eady8382 (2026).

30. Wahle, B.M., et al. Chemopreventative celecoxib fails to prevent schwannoma formation or sensorineural hearing loss in genetically engineered murine model of neurofibromatosis type 2. Oncotarget 9, 718–725 (2018).

31. Fuse, M.A., et al. Preclinical assessment of MEK1/2 inhibitors for neurofibromatosis type 2-associated schwannomas reveals differences in efficacy and drug resistance development. Neuro Oncol 21, 486–497 (2019).

32. Stemmer-Rachamimov, A.O., et al. Loss of the NF2 gene and merlin occur by the tumorlet stage of schwannoma development in neurofibromatosis 2. J Neuropathol Exp Neurol 57, 1164–1167 (1998).

33. Dewan, R., et al. Evidence of polyclonality in neurofibromatosis type 2-associated multilobulated vestibular schwannomas. Neuro Oncol 17, 566–573 (2015).

34. de Vries, M., et al. Tumor-associated macrophages are related to volumetric growth of vestibular schwannomas. Otol Neurotol 34, 347–352 (2013).

35. Malange, K.F., et al. Macrophages and glial cells: Innate immune drivers of inflammatory arthritic pain perception from peripheral joints to the central nervous system. Front Pain Res (Lausanne) 3, 1018800 (2022).

36. Schulz, A., et al. The importance of nerve microenvironment for schwannoma development. Acta Neuropathol 132, 289–307 (2016).

37. Zigmond, R.E. & Echevarria, F.D. Macrophage biology in the peripheral nervous system after injury. Prog Neurobiol 173, 102–121 (2019).

38. Baruah, P., et al. Single-cell RNA sequencing analysis of vestibular schwannoma reveals functionally distinct macrophage subsets. Br J Cancer 130, 1659–1669 (2024).

39. Yidian, C., Chen, L., Hongxia, D., Yanguo, L. & Zhisen, S. Single-cell sequencing reveals the cell map and transcriptional network of sporadic vestibular schwannoma. Front Mol Neurosci 15, 984529 (2022).

40. Feng, R., Muraleedharan Saraswathy, V., Mokalled, M.H. & Cavalli, V. Self-renewing macrophages in dorsal root ganglia contribute to promote nerve regeneration. Proc Natl Acad Sci U S A 120, e2215906120 (2023).

41. Heller, B.A., et al. Functionally distinct PI 3-kinase pathways regulate myelination in the peripheral nervous system. J Cell Biol 204, 1219–1236 (2014).

42. Doh, H.M., Kozlova, N., Cruz, K.A. & Muranen, T. Interaction of NDRG1 and TGM2 modulates DNA replication and repair. Mol Cancer Res (2026).

43. Etuze, T., Repesse, Y., Vivien, D. & Dubois, F. NDRG1 as a sensor and mediator of endothelial stress: from homeostasis to inflammation and cardiovascular disease. Cell Mol Life Sci 82, 402 (2025).

44. Kozlova, N., et al. Cancer-associated fibroblasts regulate DNA repair in pancreatic cancer through NDRG1-mediated R-loop processing. Nat Cell Biol 28, 986–999 (2026).

45. Doh, H.M., et al. Interaction of NDRG1 and MRE11 Modulates DNA Replication and Repair. Cancers (Basel) 18(2026).

46. Collinet, C., et al. Systems survey of endocytosis by multiparametric image analysis. Nature 464, 243–249 (2010).

47. James, M.F., et al. NF2/merlin is a novel negative regulator of mTOR complex 1, and activation of mTORC1 is associated with meningioma and schwannoma growth. Mol Cell Biol 29, 4250–4261 (2009).

48. Plotkin, S.R. & Wick, A. Neurofibromatosis and Schwannomatosis. Semin Neurol 38, 73–85 (2018).

49. Stierli, S., Imperatore, V. & Lloyd, A.C. Schwann cell plasticity-roles in tissue homeostasis, regeneration, and disease. Glia 67, 2203–2215 (2019).

50. Jones, A.P., et al. Spatial mapping of immune cell environments in NF2-related schwannomatosis vestibular schwannoma. Nat Commun 16, 2944 (2025).

51. Gregory, G.E., et al. Alternatively activated macrophages are associated with faster growth rate in vestibular schwannoma. Brain Commun 6, fcae400 (2024).

52. Liu, P., et al. Role of macrophages in peripheral nerve injury and repair. Neural Regen Res 14, 1335–1342 (2019).

53. Cao, J. & Liu, C. Mechanistic studies of tumor-associated macrophage immunotherapy. Front Immunol 15, 1476565 (2024).

54. Tirosh, I. & Suva, M.L. Cancer cell states: Lessons from ten years of single-cell RNA-sequencing of human tumors. Cancer Cell 42, 1497–1506 (2024).

55. Schonkeren, S.L., et al. Nervous NDRGs: the N-myc downstream-regulated gene family in the central and peripheral nervous system. Neurogenetics 20, 173–186 (2019).

56. McClatchey, A.I., et al. Mice heterozygous for a mutation at the Nf2 tumor suppressor locus develop a range of highly metastatic tumors. Genes Dev 12, 1121–1133 (1998).

57. Ruiz-Torres, D.A., et al. Spatial characterization of tertiary lymphoid structures as predictive biomarkers for immune checkpoint blockade in head and neck squamous cell carcinoma. Oncoimmunology 14, 2466308 (2025).

