## Supplemental Information for "Quantitative imaging of schwannoma captures heterogeneity and accelerates preclinical testing, exposing distinct therapeutic signatures"

Wright et al. Supplemental Figure 1

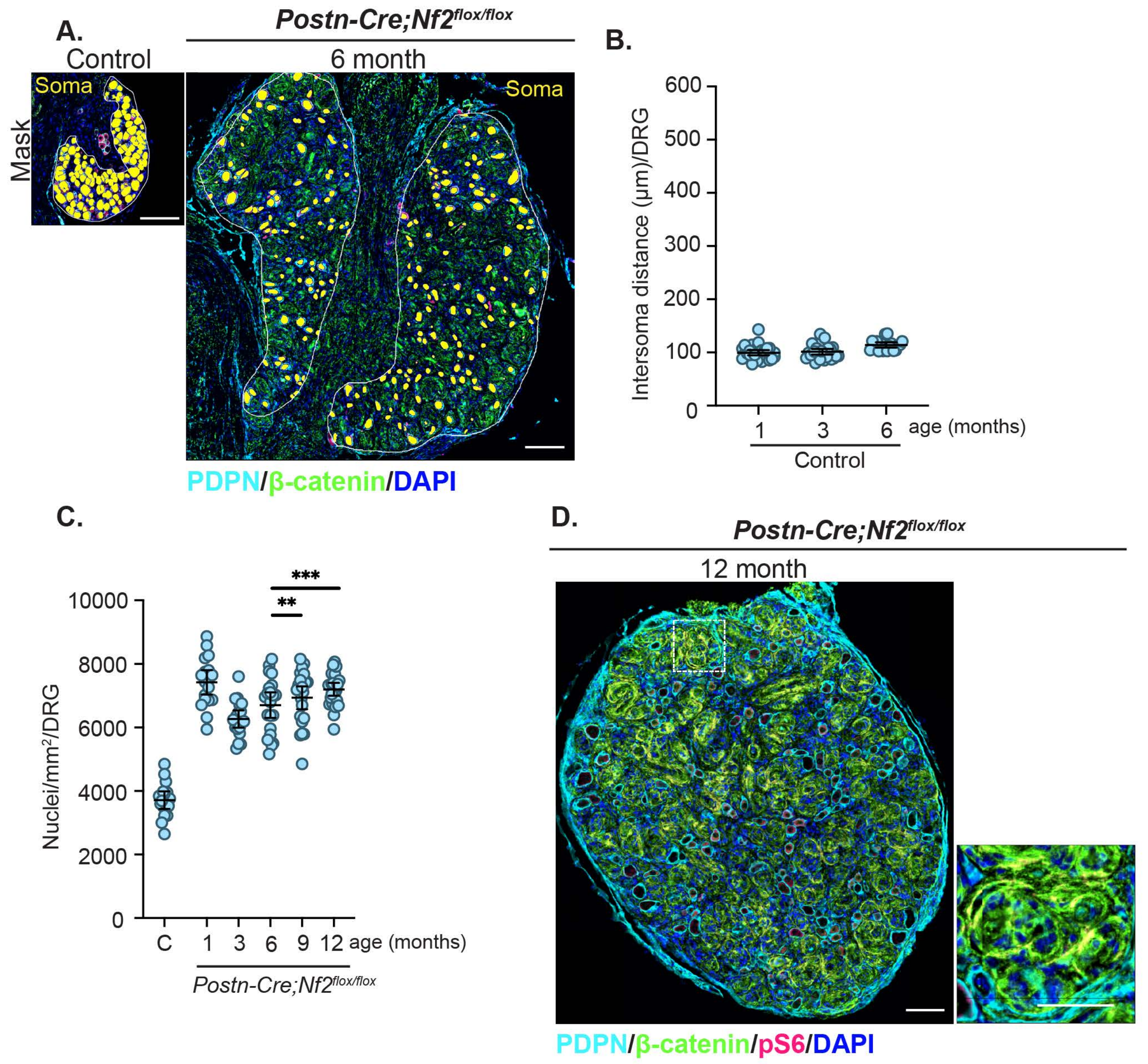

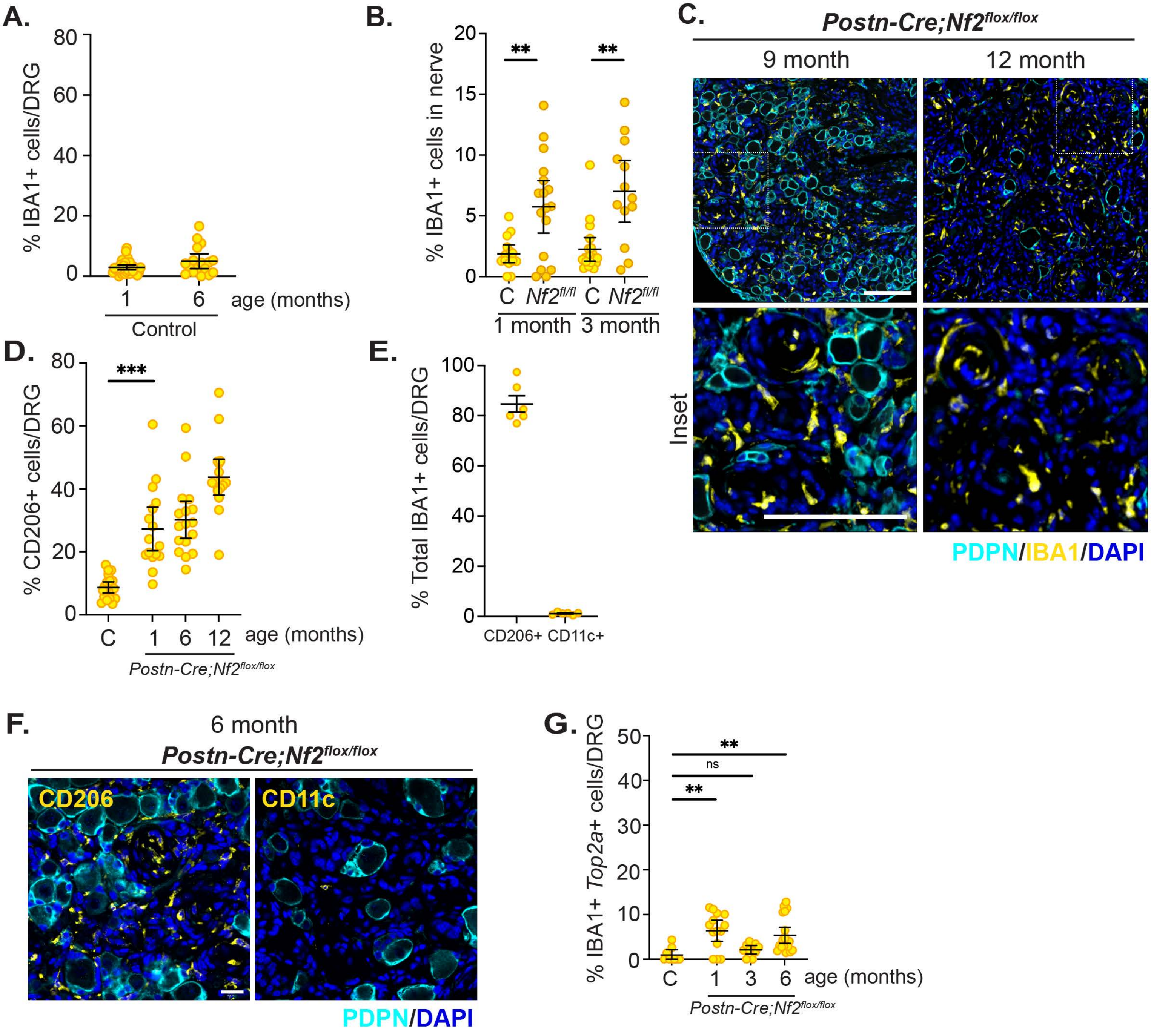
A.
% IBA1+ cells/DRG
80
60
40
20
0
1
6
age (months)
Control
B.
% IBA1+ cells in nerve
20
15
10
5
0
**
**
C
Nf2fl/fl
C
Nf2fl/fl
1 month
3 month
C.
Postn-Cre;Nf2flox/flox
9 month
12 month
Inset
PDPN/IBA1/DAPI
D.
% CD206+ cells/DRG
80
60
40
20
0
***
C
1
6
12
age (months)
Postn-Cre;Nf2flox/flox
E.
% Total IBA1+ cells/DRG
100
80
60
40
20
0
CD206+
CD11c+
F.
6 month
Postn-Cre;Nf2flox/flox
CD206
CD11c
PDPN/DAPI
G.
% IBA1+ Top2a+ cells/DRG
50
40
30
20
10
0
**
ns
**
C
1
3
6
age (months)
Postn-Cre;Nf2flox/flox

Wright et al. Supplemental Figure 3

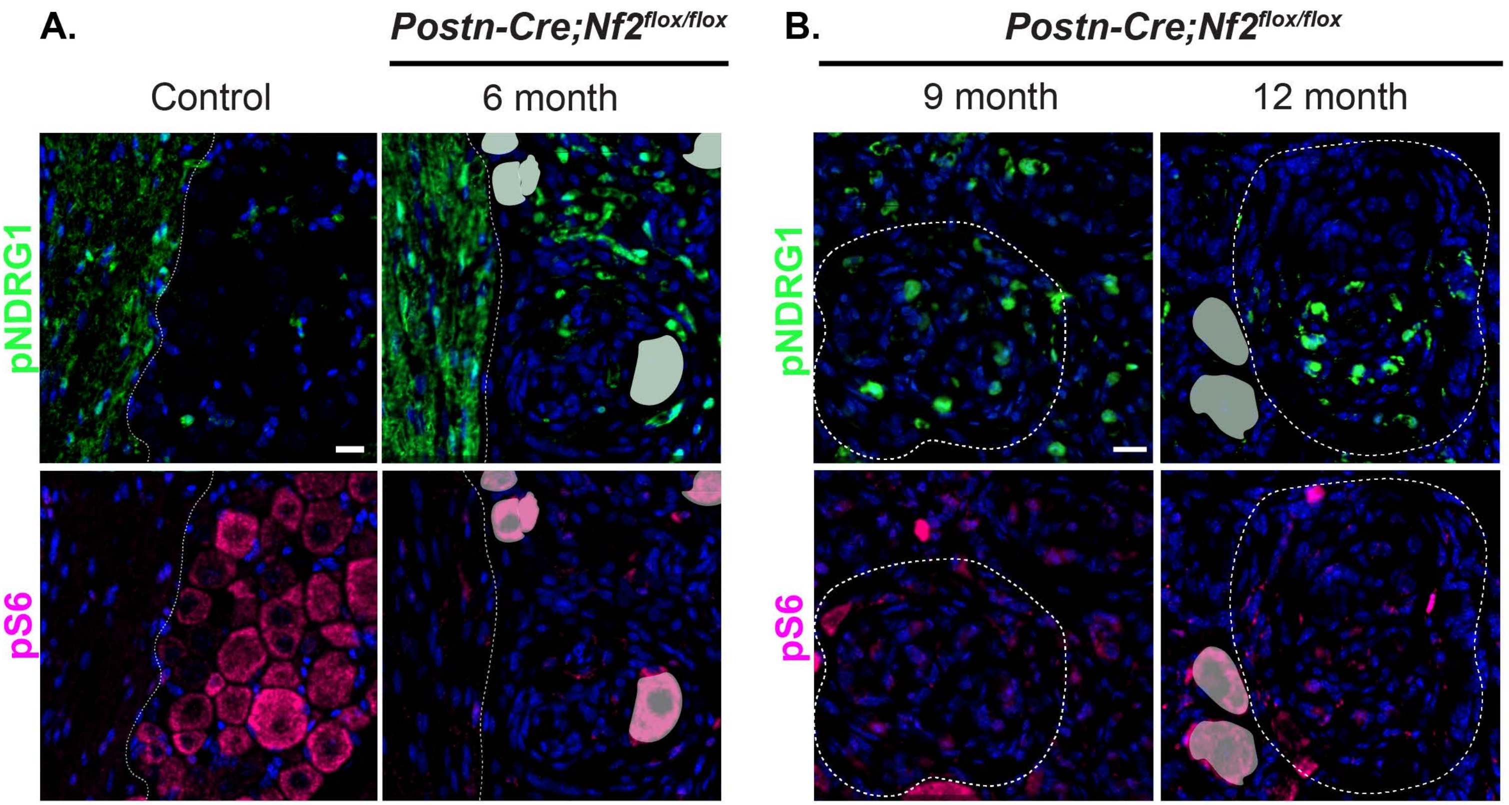

Wright et al. Supplemental Figure 4

**A.**

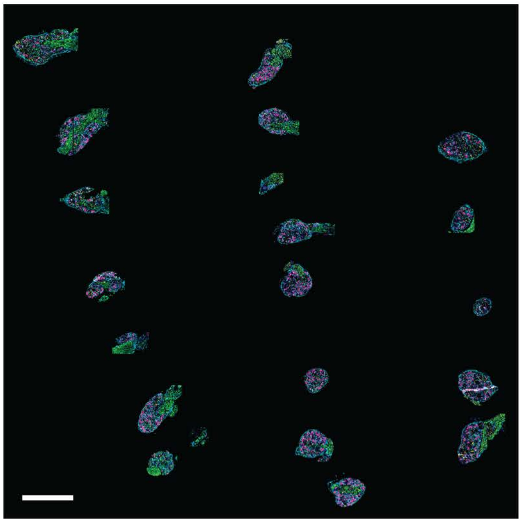

Wright et al. Supplemental Figure 5

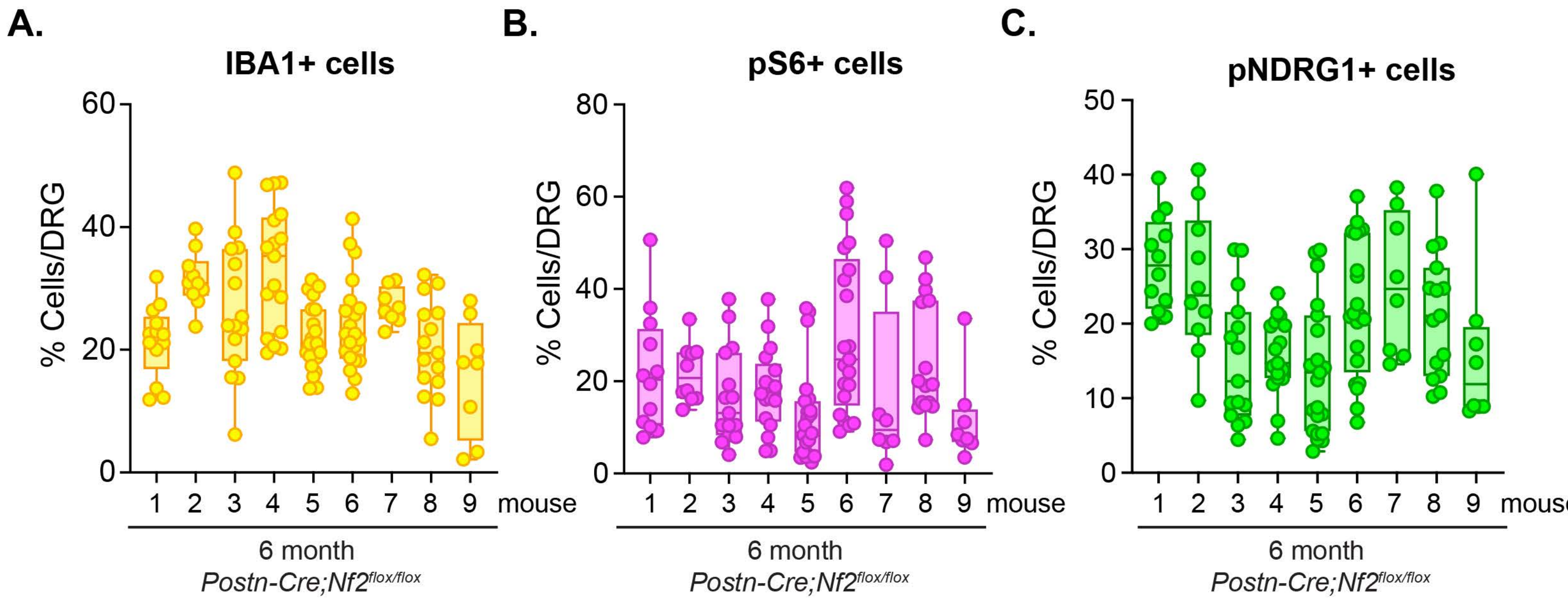

Wright et al. Supplemental Figure 6

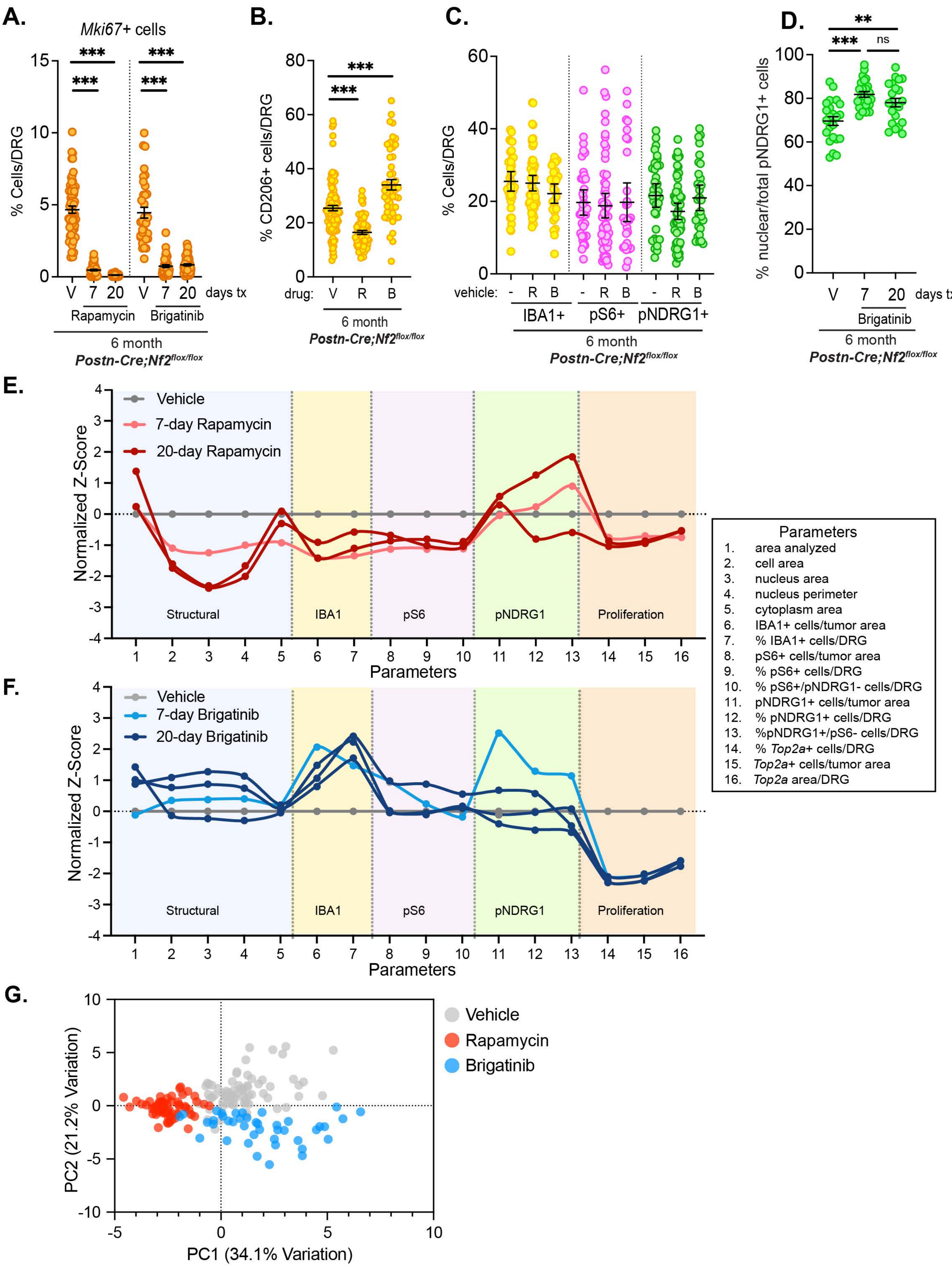

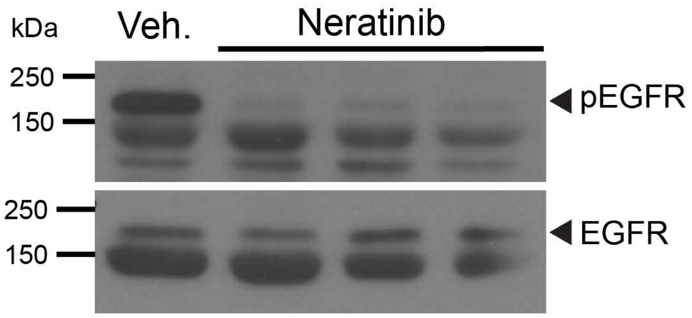
kDa
Veh.
Neratinib
250
150
pEGFR
250
150
EGFR

**Supplemental Figure legends**

**Figure S1. Related to Figure 1.** A. Single 6 month-old control *Nf2*$^{flox/flox}$ (left) and *Postn-Cre;Nf2*$^{flox/flox}$ (right) DRG as examples of HALO soma segmentation masks (yellow) used for measuring intersoma distance (PDPN, cyan; β-catenin, green). Note that the DRG are shown at the same scale to highlight the much larger mutant DRG. Scale bar = 100 µm. B. Quantitation of intersoma distance in control DRG from 1, 3, and 6 month-old mice. Each datapoint represents a single DRG. C. Quantitation of the density of nuclei in the abnormal interstitial tissue in *Postn-Cre;Nf2*$^{flox/flox}$ DRG over time. Note that the small drop in nuclear density at 3 months corresponds with the clear increase in membrane area labelled by β-catenin in the main figure (Fig 1A, insets). Bars represent the mean +/- 95% CI. (**p<0.01, ***p<0.001). D. Scaled images of representative 12 month-old *Postn-Cre;Nf2*$^{flox/flox}$ DRG stained with antibodies that detect SGCs (anti-podoplanin PDPN, cyan) surrounding each neuronal soma, membrane (anti-β-catenin, green) and neuronal soma (anti-pS6; magenta). Nuclei are labeled with DAPI (blue). Inset highlights 'whorls' that begin to appear at 3 months and expand over time. Note that pS6 also marks some supernumerary interstitial tumor cells. Scale bars: 100 µm main, 50 µm inset.

**Figure S2. Related to Figure 2**. A. Quantitation of IBA1+ macrophages in control DRG at 1 and 6 months of age (PDPN+ SGCs, cyan). B. Quantitation of IBA1+ macrophages in the nerve root of 1 and 3 month-old control (C) and *Postn-Cre;Nf2*$^{flox/flox}$ DRG. C. *Top,* Representative immunofluorescent images of macrophages (Iba1+, yellow) in DRG from 9 and 12 month-old *Postn-Cre;Nf2*$^{flox/flox}$ mice. *Bottom*, magnified insets showing extensive interactions of macrophages with aberrant SGCs (PDPN+, cyan) and interstitial cells. Nuclei are labeled with DAPI (blue). Scale bars = 100 µm. D. Quantitation of CD206+ cells in control and *Postn-Cre;Nf2*$^{flox/flox}$ DRG at 1, 6, and 12 months of age; compare to quantitation of IBA+ cells in Figure 2B. E. Quantitation of Iba1+ cells in 6 month *Postn-Cre;Nf2*$^{flox/flox}$ DRG that co-express CD206 or CD11c. F. Representative images of CD206+ (left, yellow) and CD11c+ (right, yellow) macrophages in 6 month-old *Postn-Cre;Nf2*$^{flox/flox}$ DRG (PDPN+ SGCs, cyan). Nuclei are labeled with DAPI (blue). Scale bars = 20 µm. G. Quantitation of IBA1+ macrophages that express the proliferation marker *Top2a* as detected by RNAscope. Note that the number of *Top2a*+ macrophages in the *Postn-Cre;Nf2*$^{flox/flox}$ 1 month DRG, which is slightly increased over the control, corresponds to 16/mm$^2$, which is vastly lower than what has been quantified in the DRG post-injury by others (4000/mm$^2$)[40]. Each datapoint represents one DRG. Bars represent mean +/- 95% CI. (**p<0.01, ***p<0.001, ns = not significant).

**Figure S3. Related to Figure 3.** A. Representative images of control and *Postn-Cre;Nf2$^{flox/flox}$* DRG showing a portion of the nerve containing myelinating Schwann cells that stain strongly for membrane localized pNDRG1 (green, top) but are largely devoid of pS6 (magenta, bottom). The nerve root is to the left of the dotted line. Note that pNDRG1 in the abnormal interstitial tissue is often nuclear. Soma are masked in the mutant as in Figure 3. B. Representative images of DRG from 9 and 12 month-old *Postn-Cre;Nf2$^{flox/flox}$* mice stained with pNDRG1 (green, top) and pS6 (magenta, bottom). Dotted line demarcates an individual whorl in each image. Scale bars = 20 µm.

**Figure S4. Related to Figure 4.** A. Representative whole-slide scan of an array of 20 DRG from a 6 month old *Postn-Cre;Nf2$^{flox/flox}$* mouse stained for PDPN (cyan), Iba1 (yellow), pS6 (magenta) and pNDRG1 (green). Scale bars = 1 mm.

**Figure S5. Related to Figure 5.** A. Quantitation of Iba1+ cells in multiple DRG from 9 different 6 month old *Postn-Cre;Nf2$^{flox/flox}$* mice. B. Quantitation of pS6+ cells in multiple DRG from 9 different 6 month old *Postn-Cre;Nf2$^{flox/flox}$* mice. C. Quantitation of pNDRG1+ cells in multiple DRG from 9 different 6 month old *Postn-Cre;Nf2$^{flox/flox}$* mice. Each data point represents one DRG.

**Figure S6. Related to Figure 6.** A. Quantitation of *Mki67*+ cells from 6 month old *Postn-Cre;Nf2$^{flox/flox}$* mice treated for 7 or 20 days (5/7 days per week for 4 weeks) with vehicle (V), rapamycin (R, 8 mg/kg) or brigatinib (B, 50 mg/kg). B. Quantitation of CD206+ cells from 6 month-old *Postn-Cre;Nf2$^{flox/flox}$* mice treated for 7 days with vehicle (V), rapamycin (R) or brigatinib (B); compare to the quantitation of IBA1 in Figure 6A. C. Comparative quantitation of IBA1+, pS6+, and pNDRG1+ cells from 6 month-old *Postn-Cre;Nf2$^{flox/flox}$* mice untreated or treated for 7 days with vehicles for Rapamycin (R) or Brigatinib (B). D. Quantitation of the percentage of pNDRG1+ cells that exhibit nuclear pNDRG1 localization in 6 month-old *Postn-Cre;Nf2$^{flox/flox}$*

mice treated for 7 or 20 days with vehicle (V) or brigatinib (B). For A-D, each datapoint represents one DRG. Bars represent the mean +/- 95% CI. (***p<0.001) E. Multiparametric graph assembling 16 different parameters measured using the HighPlex FL module in HALO across mice treated with vehicle (green, average of 3 mice) rapamycin for 7 (gray, average of 3 mice) or 20 days (red, each line represents one mouse). Graphs were plotted as described in Figure 6. F. Multiparametric graph assembling 16 different parameters measured using the HighPlex FL module in HALO across mice treated with vehicle (green, average of 3 mice) brigatinib for 7 (gray, average of 3 mice) or 20 days (blue, each line represents one mouse). Multiparametric graphs were plotted as described in Figure 6. G. PCA plot of the parameters measured in figure 6D in vehicle-, rapamycin-, or brigatinib-treated mice. Each dot represents one DRG.

**Figure S7. Related to Figure 7.** Immunoblot showing levels of phospho- and total- EGFR in mice treated for 7 days with vehicle or neratinib.
